# Visual pigment evolution in Characiformes: the dynamic interplay of teleost whole-genome duplication, surviving opsins and spectral tuning

**DOI:** 10.1101/695544

**Authors:** Daniel Escobar-Camacho, Karen L. Carleton, Devika W. Narain, Michele E.R. Pierotti

## Abstract

Vision represents an excellent model for studying adaptation, given the genotype-to-phenotype-map that has been characterized in a number of taxa. Fish possess a diverse range of visual sensitivities and adaptations to underwater light making them an excellent group to study visual system evolution. In particular, some speciose but understudied lineages can provide a unique opportunity to better understand aspects of visual system evolution such as opsin gene duplication and neofunctionalization. In this study, we characterized the visual system of Neotropical Characiformes, which is the result of several spectral tuning mechanisms acting in concert including gene duplications and losses, gene conversion, opsin amino acid sequence and expression variation, and A_1_/A_2_-chromophore shifts. The Characiforms we studied utilize three cone opsin classes (SWS2, RH2, LWS) and a rod opsin (RH1). However, the characiform’s entire opsin gene repertoire is a product of dynamic evolution by opsin gene loss (SWS1, RH2) and duplication (LWS, RH1). The LWS- and RH1-duplicates originated from a teleost specific whole-genome duplication as well as characiform-specific duplication events. Both LWS-opsins exhibit gene conversion and, through substitutions in key tuning sites, one of the LWS-paralogs has acquired spectral sensitivity to green light. These sequence changes suggest reversion and parallel evolution of key tuning sites. In addition, characiforms exhibited species-specific differences in opsin expression. Finally, we found interspecific and intraspecific variation in the use of A_1_/A_2_-chromophores correlating with the light environment. These multiple mechanisms may be a result of the highly diverse visual environments where Characiformes have evolved.

## Introduction

To fully understand the evolutionary history of genes and their relevance for adaptation and speciation, it is important to explore the genotype to phenotype map and how this relates to the environment. Evolutionary studies of genes such as the ones involved in the first steps of vision can provide valuable insights in the acquisition of new functions and their adaptive significance. In vertebrates, vision starts when light reaches the retina and is detected by rod (night vision) or cone (diurnal vision) photoreceptors. Photoreceptors are packed with visual pigments that are composed of two components: an opsin protein with seven α-helices enclosing a ligand-binding pocket, and a light-sensitive chromophore, 11-cis retinal (Bowmaker 2008; Yokoyama 2008). There can be multiple cone types containing different visual pigments that absorb light maximally in different parts of the wavelength spectrum.

There are four classes of cone pigments encoded by opsin genes among vertebrates: a short-wave class (SWS1) sensitive to ultraviolet-violet light (350-400 nm), a second short-wave class (SWS2) sensitive to violet-blue (410-490 nm), a middle-wave class (RH2) sensitive to green (480-535 nm), and a middle- to long-wave class (LWS) sensitive to the green and red spectral region (490-570 nm) (Bowmaker and Hunt 2006; Bowmaker 2008). All four cone classes are the product of a series of gene duplications from an ancestral single opsin gene that appeared early in vertebrate evolution (450 MYA) (Bowmaker 1998; Bowmaker and Hunt 2006; Bowmaker 2008). This results in a spectral tuning mechanism that is based on nucleotide variation: if a nucleotide substitution leads to the replacement of an amino acid that alters the interaction of the chromophore and the opsin, this will lead to a spectral shift in the maximal absorbance (λ_max_) of the visual pigment. Consequently, variation in λ_max_ between visual pigments is the product of the interaction of different opsin classes and the identical 11-cis retinal. The shift in λ_max_ caused by a single amino acid substitution depends on the amino acid identity and site, with most causing smaller shifts (2-10 nm) and a few sites causing very large shifts (e.g. 75 nm) (Yokoyama 2008).

Among vertebrates, fish are ideal for the study of visual pigment evolution. First, because of the physico-chemical properties of water, this medium has a profound effect on light transmission. Water absorbs and scatters much of the incoming light, and this inevitably causes great variation across aquatic habitats that differ in concentrations of particulates and dissolved compounds (Loew and McFarland 1990; Warrant and Johnsen 2013). This results in several adaptations in fish visual systems. Second, due to their phylogenetic history, species richness, diverse ecologies, and diverse spectral sensitivities, teleosts offer an excellent system for studying the evolution of visual pigments. Spectral sensitivities have been documented for quite a few fish species (Schwanzara 1967; Muntz 1973; Levine and MacNichol 1979; Bowmaker et al. 1994; Lythgoe et al. 1994; Carleton 2009) and the dynamic evolution of the different opsin classes has been actively studied (Bowmaker 2008; Yokoyama 2008; Hofmann and Carleton 2009; Davies et al. 2012; Rennison et al. 2012; Cortesi et al. 2015; Lin et al. 2017; Musilova et al. 2019).

Characiformes, with more than 2000 described species, is an extremely diverse group of freshwater fishes inhabiting a wide range of ecosystems. This order includes at least 23 families with dozens of species being described each year (Oliveira et al. 2011; Arcila et al. 2018; Froese and Pauly 2019). Their Gondwanan origin, wide distribution, species richness and colorful patterns, make them an ideal group for studying the evolution of their visual system and its adaptation to the light environment. Data on the opsin repertoire of Characiformes has only been reported for the tetra *Astyanax fasciatus* (Yokoyama and Yokoyama 1990a; Yokoyama and Yokoyama 1993; Register et al. 1994; Yokoyama et al. 1995; Yokoyama et al. 2008). These studies characterized its visual pigments and showed how this species has a duplication in the LWS opsin in which one copy became sensitive to green light through amino acid substitutions; a remarkable example of convergent evolution with green sensitivity in humans (Yokoyama and Yokoyama 1990a). Recent studies have analyzed the origins of *Astyanax* opsin genes more in depth and concluded that these duplicates are surviving opsins from the teleost-specific genome duplication (TGD) (300-450 MYA: Taylor et al. 2001; Meyer and Peer 2005; Liu et al. 2018).

In this study, we expand the molecular characterization of the visual system in Characiformes. We showcase the complex evolutionary dynamics of their opsin gene repertoire and we examine the diverse set of spectral tuning mechanisms present in this group.

## Results

### Opsin gene sequences

#### Opsin complements

Through phylogenetic analyses of 15 characiform species, we identified fully functional sequences belonging to three cone opsin classes (SWS2, RH2, LWS) as well as the rod opsin (RH1) (Fig. 1-2, Fig S1-3). The opsin-gene set within Characiformes seems highly variable because we found variation in the presence/absence of some opsins as well as several duplications (Fig. 3). We did not find sequences belonging to the UV-light sensitive opsin (SWS1), either in the transcriptomes or genomes (Fig. S1). We also did not detect the RH2 opsin in the transcriptomes of *C. spilurus, H. microlepis, S. rhombeus*, and *R. guatemalensis*; however, we found a non-functional RH2 opsin in the genome of *P. nattereri*.

**Figure 1.**
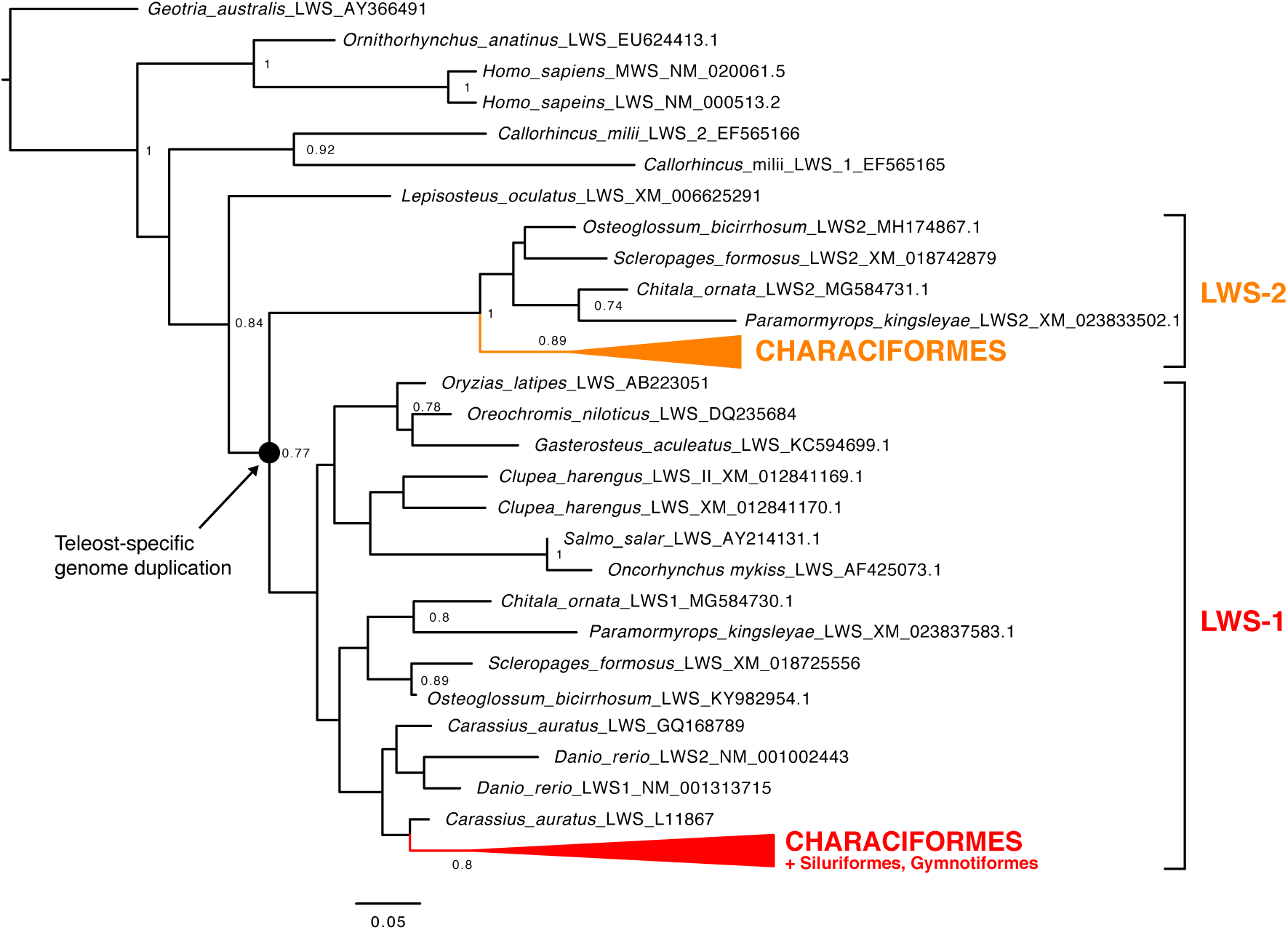
LWS opsin tree of Characiformes. LWS opsin maximum-likelihood phylogenetic tree based on amino-acid sequences of Characiformes, Osteoglossiformes, Siluriformes, Gymnotiformes, *Geotria australis* (lamprey), *Ornithorhynchus anatinus* (platypus), *Homo sapiens* (humans), *Callorhinchus milii* (Elephant shark), *Lepisosteus oculatus* (Spotted gar), *Oryzias latipes* (medaka), *Gasterosteus aculeatus* (stickleback), *Clupea harengus* (herring), *Salmo salar* (salmon), *Oncorhynchus mykiss* (trout), *Carassius auratus* (goldfish), and *Danio rerio* (zebrafish). Bootstrap support over 75% is shown. This tree confirms that LWS1 and LWS2 arose after the divergence of the spotted gar, probably as a product of teleost whole genome duplication (TGD). Notice the clustering of characiform LWS2 opsins with the osteoglossimorph LWS2 opsins. Characiform species are represented as compressed color-filled clades (*LWS2 in orange and LWS1 in red*).

**Figure 2.**
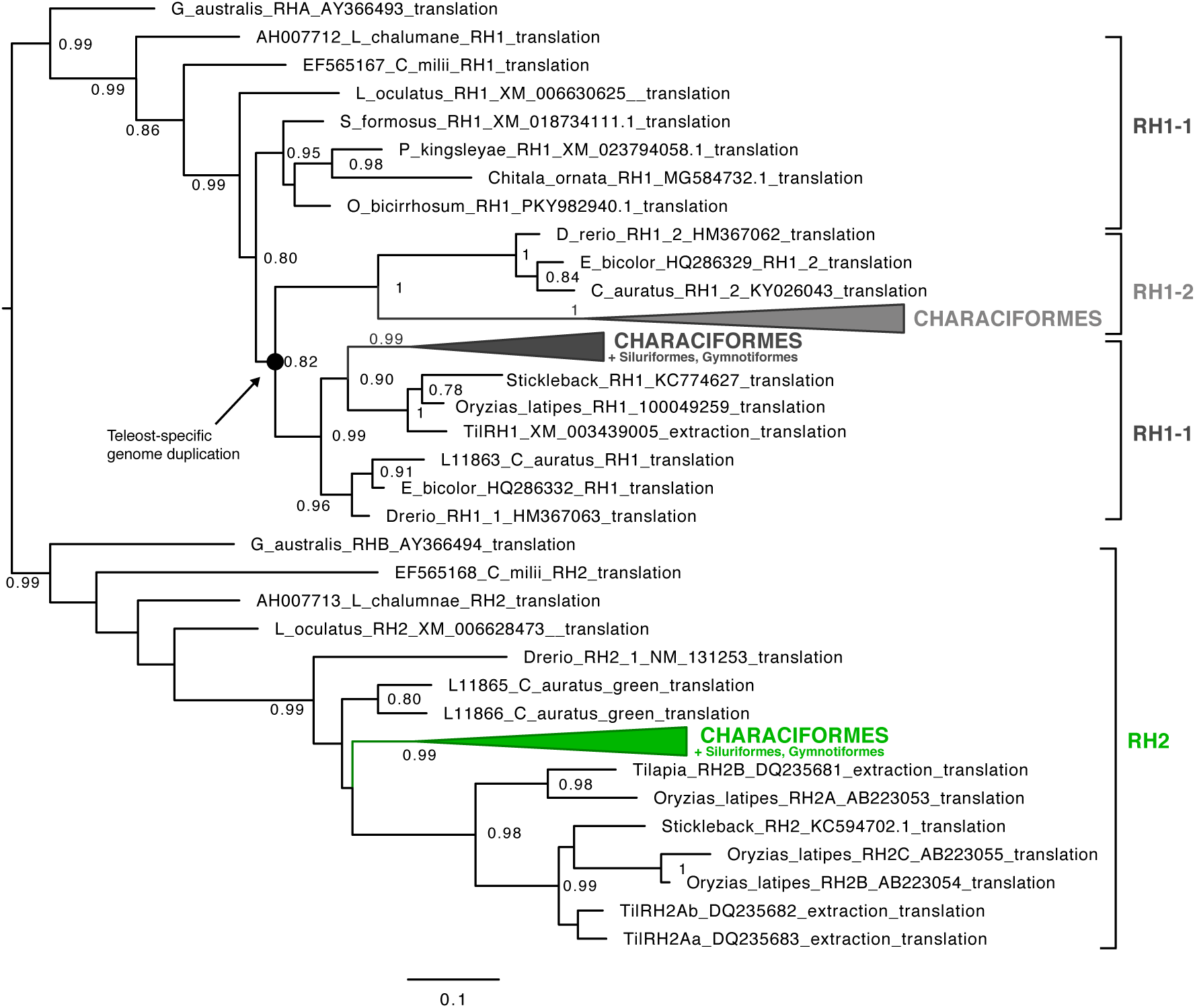
RH1-RH2 opsin tree of Characiformes. RH1 and RH2 opsin maximum-likelihood phylogenetic tree based on amino-acid sequences of Characiformes, Osteoglossiformes, Siluriformes, Gymnotiformes, Cypriniformes, *Geotria australis* (lamprey), *Latimeria calumnae* (coelacant), *Callorhinchus milii* (Elephant shark), *Lepisosteus oculatus* (Spotted gar), *Oryzias latipes* (medaka), and *Gasterosteus aculeatus* (stickleback). Bootstrap support over 75% is shown. This tree confirms that RH1-2 arose after the divergence of the spotted gar, probably as a product of teleost whole genome duplication (TGD). Notice the clustering of characiform RH1-2 opsins with the cyprinimorphs surviving RH1-2 opsins. Characiform species are compressed in color-filled clades (*RH1-2 in gray, RH1-1 in black, and RH2 in green*).

**Figure 3.**
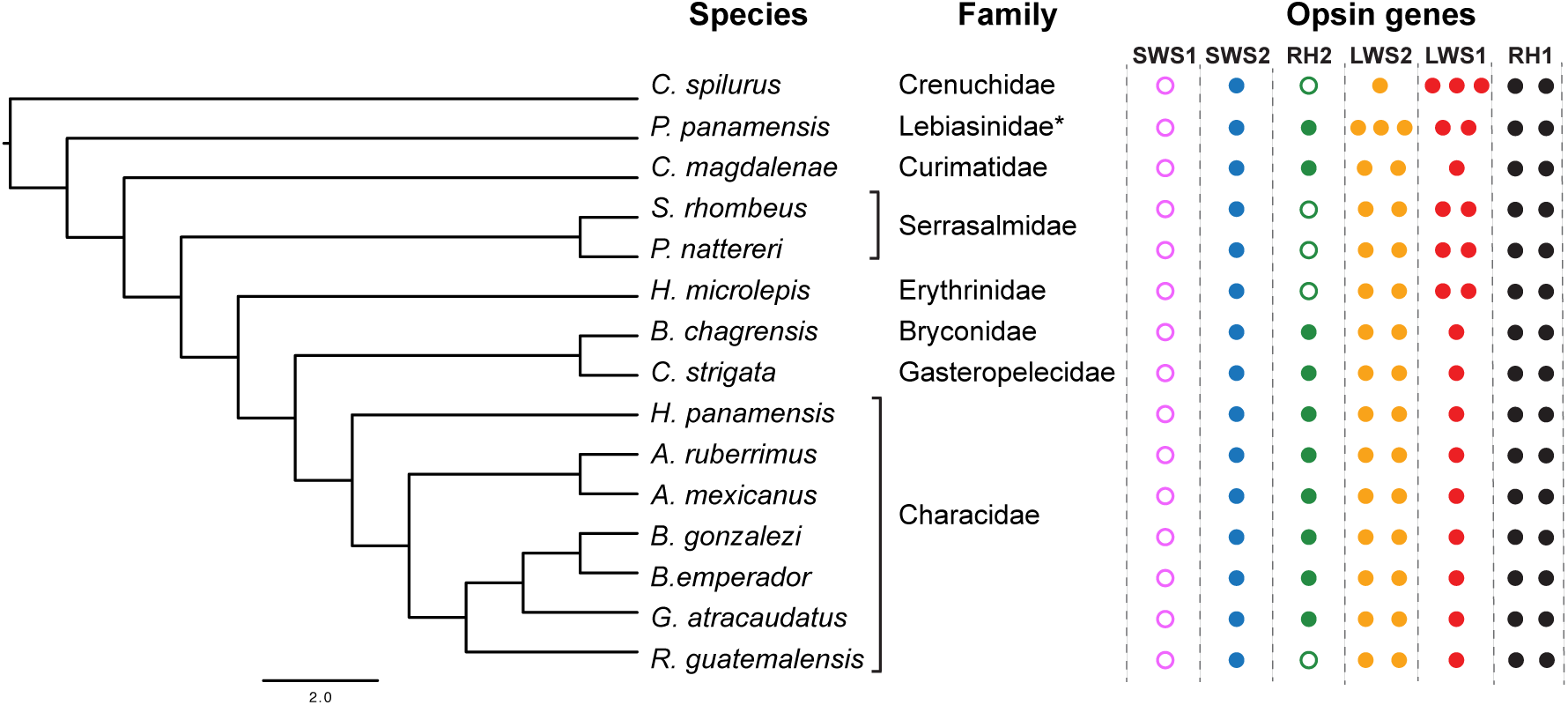
Opsin gene complement in Characiformes. To the left, schematic representation of the phylogenetic relationships of characiforms in this study based on Oliveira et al. (2011), and to the right, based on transcriptomes and genomes, the presence or absence as well as the number of opsin genes in each class. Species names and families are shown for the samples used in this study as well as their opsin complement where each opsin-gene is indicated by a filled circle for each opsin class. Empty circles denote potential gene losses. *Even though *P. panamensis* is considered a member of Lebiasinidae, our results grouped this species with the Parodontidae.

Examination of the LWS opsin class revealed the presence of several duplications and these varied between lineages suggesting the following. First, there was an LWS-duplication event, which was the product of the teleost-specific genome duplication (TGD) (300-450 MYA) (Fig. S4). This is supported by the fact that these initial duplicates appear after the divergence of teleosts and the spotted gar, *Lepisosteus oculatus* (Fig. 1). These initial LWS paralogs formed two distinct clades in our analyses, which we will call LWS1 and LWS2. LWS2 opsins grouped with Osteoglossiformes, which are known to share this duplication with characiforms (Liu et al. 2018), whereas the LWS1 opsins grouped with the remaining teleost LWS opsins. Second, after TGD, LWS1 and LWS2 underwent subsequent rounds of duplications within Characiformes. This varied across families ranging from having one to up to three LWS-duplicates (Fig. S2). The characiform unique LWS2 opsin duplication is shared in most species. LWS2-1 and LWS2-2 differ by the presence of a 6bp deletion in the first 20 bp of the coding sequence (exon I, extracellular region) in LWS2-2, and by a few amino acids, although not in spectral tuning sites.

Furthermore, we found TGD-surviving duplicates in the rodopsin (RH1) (Fig. 2). In similar fashion to the LWS duplicates, these RH1 paralogs (which we will refer to as RH1-1 and RH1-2) grouped in different RH1 clades where the TGD-surviving opsins of Characiformes formed a well-supported clade with the known TGD-surviving copies (RH1-2) present in Cypriniformes (Morrow et al. 2011; Morrow et al. 2017) (Fig. 2). Within RH1-2 opsin sequences, several species of characiforms had numerous deletions in the last exon. We also found disparate amino acid variation at transmembrane sites which suggest the non-functionality of these opsin genes, hence, they were excluded from subsequent analyses. Lastly, the LWS1, RH2, SWS2, and RH1-1 opsins of Characiformes show a paraphyletic pattern in relation to Siluriformes and Gymnotiformes (Fig. S1-3).

#### Opsin sequence spectral tuning

Our analyses revealed several amino acid substitutions that shift λ_max_ of visual pigments. The SWS2 opsin showed the greatest variation in transmembrane regions, changes in polarity, and variation in binding pocket sites (Fig. 4). Several of these substitutions occurred in spectral tuning sites (M44T, A109G, M122I, A269T, A292S) that are known to shift the SWS2 λ_max_ (Yokoyama 2008). Other opsin classes also showed variable diversity in known spectral tuning sites, including the RH2 opsin with substitutions that shift λ_max_ to shorter wavelengths (K36Q, L46F, I49C, F50L, L108T, A295S) (Chinen et al. 2005; Davies et al. 2007) and the RH1-1 opsin with substitutions that shift λ_max_ to longer wavelengths (N83D, F261Y) (Yokoyama et al. 1995; Yokoyama et al. 2005). We also confirmed the presence of mutations in three “key sites” (S164A, Y261F, T269A) in the LWS2 paralogs that shift λ_max_ to shorter wavelengths (∼30 nm) (Yokoyama 2008; Yokoyama et al. 2008). Although previously reported only in *Astyanax fasciatus* (Yokoyama and Yokoyama 1990a) and Osteoglossiformes (Liu et al. 2018), this trait appears to be present in most characiforms (Fig. 5).

**Figure 4.**
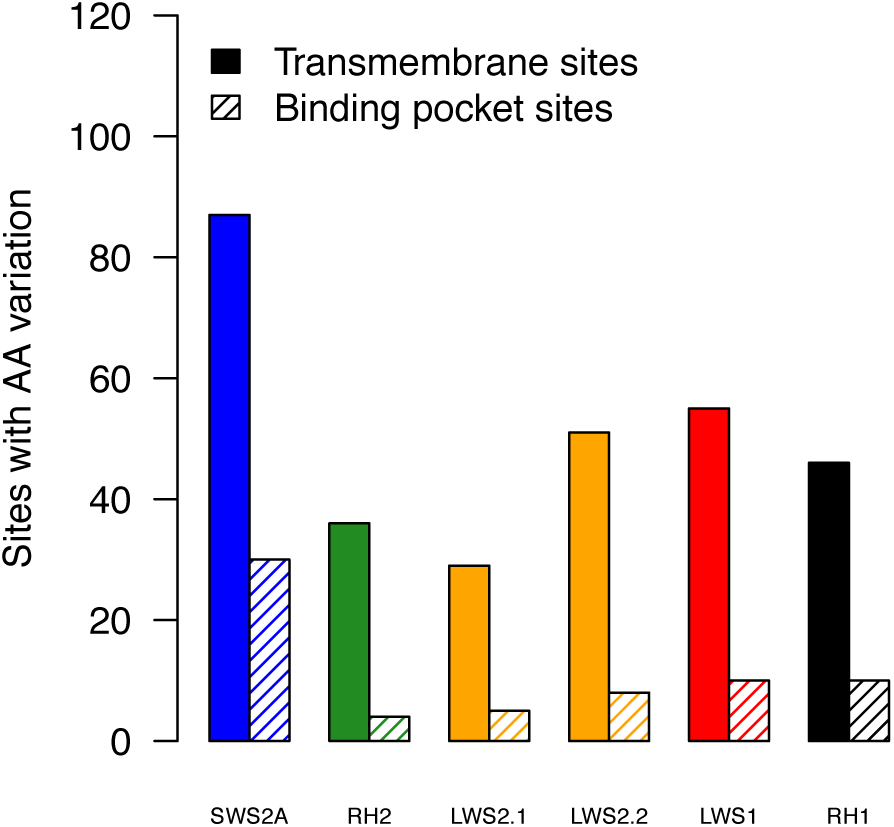
Number of sites with amino-acid substitution variation for each opsin class of 15 Characiformes species. Solid bars denote amino acid variation in transmembrane regions whereas stripped bars denote variation in binding pocket sites.

**Figure 5.**
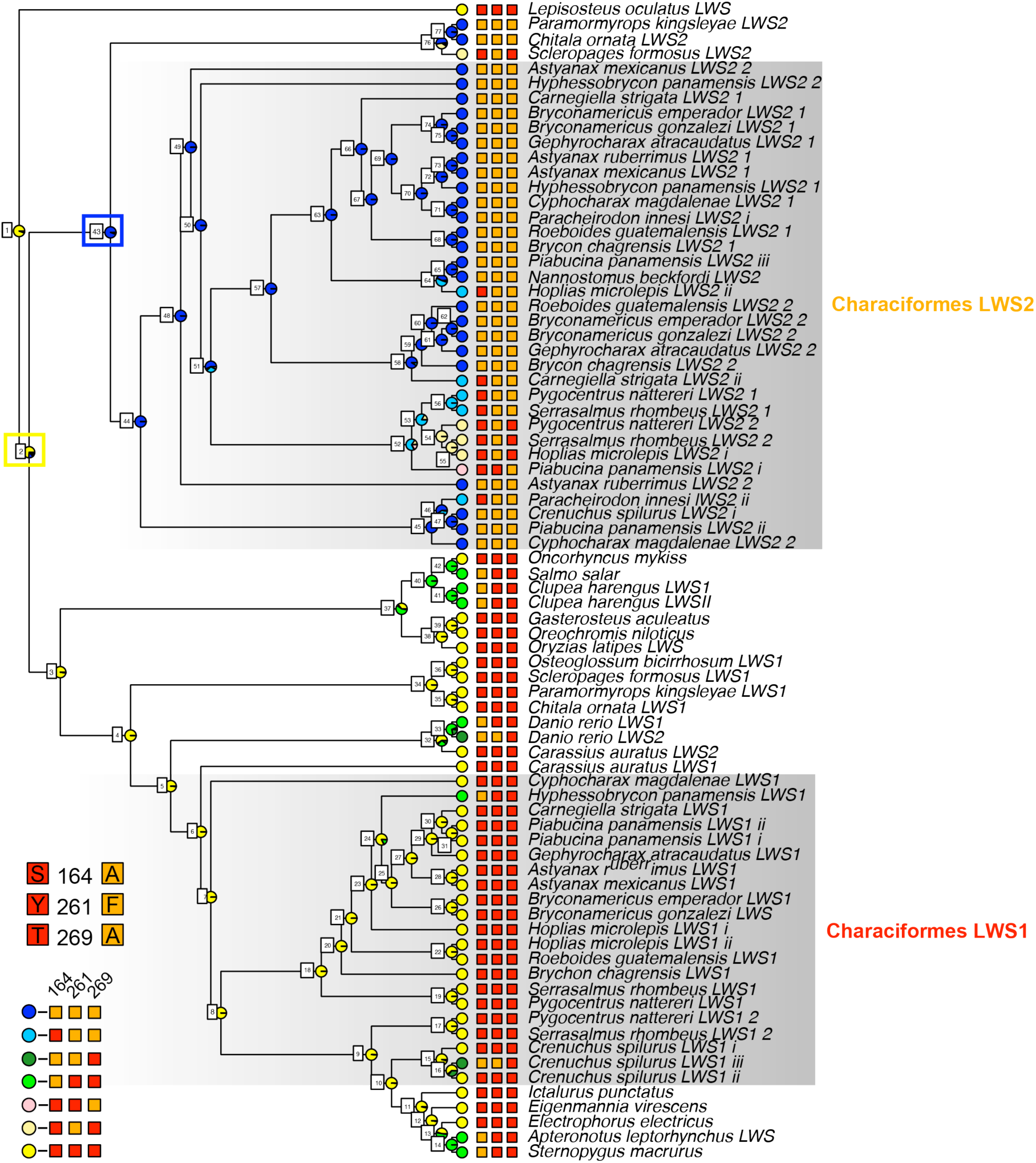
Ancestral state reconstruction of the spectral tuning in LWS opsin genes. The three known spectral tuning sites (S164A, Y261F, T269A) that are known to convey green sensitivity in Characiformes are shown for each species for each LWS gene. The seven combinations we found of the three tuning sites in our data set are also shown, with each combination represented as a colored circle. Pie charts on the nodes indicate the scaled likelihoods of each specific combination, calculated using the ace function in APE. Nodes #2 and #43 are denoted by yellow and blues rectangles respectively. Nodes are also labeled as in Table S2.

#### Gene conversion

Gene conversion analysis with GARD revealed evidence of interspecific gene conversion within LWS1 and LWS2 opsins with two and three breakpoints respectively. In both LWS1 and LWS2, conversion seems to be present in the first exons (Fig. S5, Table. S1). Phylogenetic trees based on fragments between recombination breakpoints exhibit different tree topologies. Trees based on exons three to six recovered the typical phylogenetic relationships between families and ancestral duplications based on their known genomic tree topologies (Oliveira et al. 2011) (Fig. S5).

#### Ancestral state reconstruction

By analyzing the evolution of LWS spectral tuning through ancestral state reconstruction, our results suggest that the ancestral LWS haplotype of teleosts before TGD was probably red wavelength sensitive (node #2, 73.56% of the scale likelihood) (Fig. 5, Table. S2). This suggests that green sensitivity evolved soon after TGD (node #43, 93.7% of the scaled likelihood) (Fig. 5, Table. S2).

Furthermore, our analysis examining the molecular basis of spectral tuning of the site S164A, typically conferring a −7 nm shift (Yokoyama et al. 2008; Yokoyama 2008), suggests that the LWS2 ancestral haplotype of Characiformes most probably used the codon GCC (node 43, 99.03% of the scaled likelihood) to encode for alanine whereas the LWS1-ancestral haplotype used TCT (node 6, 99% of the scaled likelihood, Table S3) to encode for serine. However, we found a reversion in the LWS2 opsins of some earlier divergent lineages within Characiformes (*C. strigata, P. nattereri, H. microlepis* and *P. panamensis*) where the reverse mutation in LWS2 opsins changed codons for alanine (GCC) back to codons for serine (TCT or TCC). This occurred in parallel in the characid *P. innesi*. (Fig. S6). Similar to LWS2, there is evidence of parallel evolution in the LWS1 opsins. *H. panamensis* and *C. spilurus* shifted in parallel from serine to alanine utilizing the same codons (TCT to GCT) (Fig. S6). Finally, even though the scope of this study focused on Characiformes, the variability of site 164 is quite extensive as several teleosts exhibit different codons for either alanine or serine (Fig. S6).

### Opsin gene expression

Opsin expression profiles varied between characiform species, ranging from some expressing mainly two opsins, like the sail-fin tetra (*C. spilurus*) or the dogfish (*H. microlepis*), to others expressing up to six (*P. panamensis*). The SWS2 opsin was the only short-wavelength pigment expressed (3 to 15% of total opsin expression), and the LWS duplicates accounted for the bulk of characiform opsin expression (80-95%) (Fig. 6). We always observed the expression of at least one copy of the LWS1 paralog, followed by the expression of one or two copies of the LWS2 paralog (Fig. 6). There seem to be differences in the expression of the LWS2 paralogs because in some species the LWS2-1 opsin is more expressed than the LWS2-2 opsin (*B. chagrensis, A. ruberrimus, G. atracaudatus*), but this pattern is reversed in other species (*H. panamensis, B. gonzalezi, C. strigata*) (Fig. 6).

**Figure 6.**
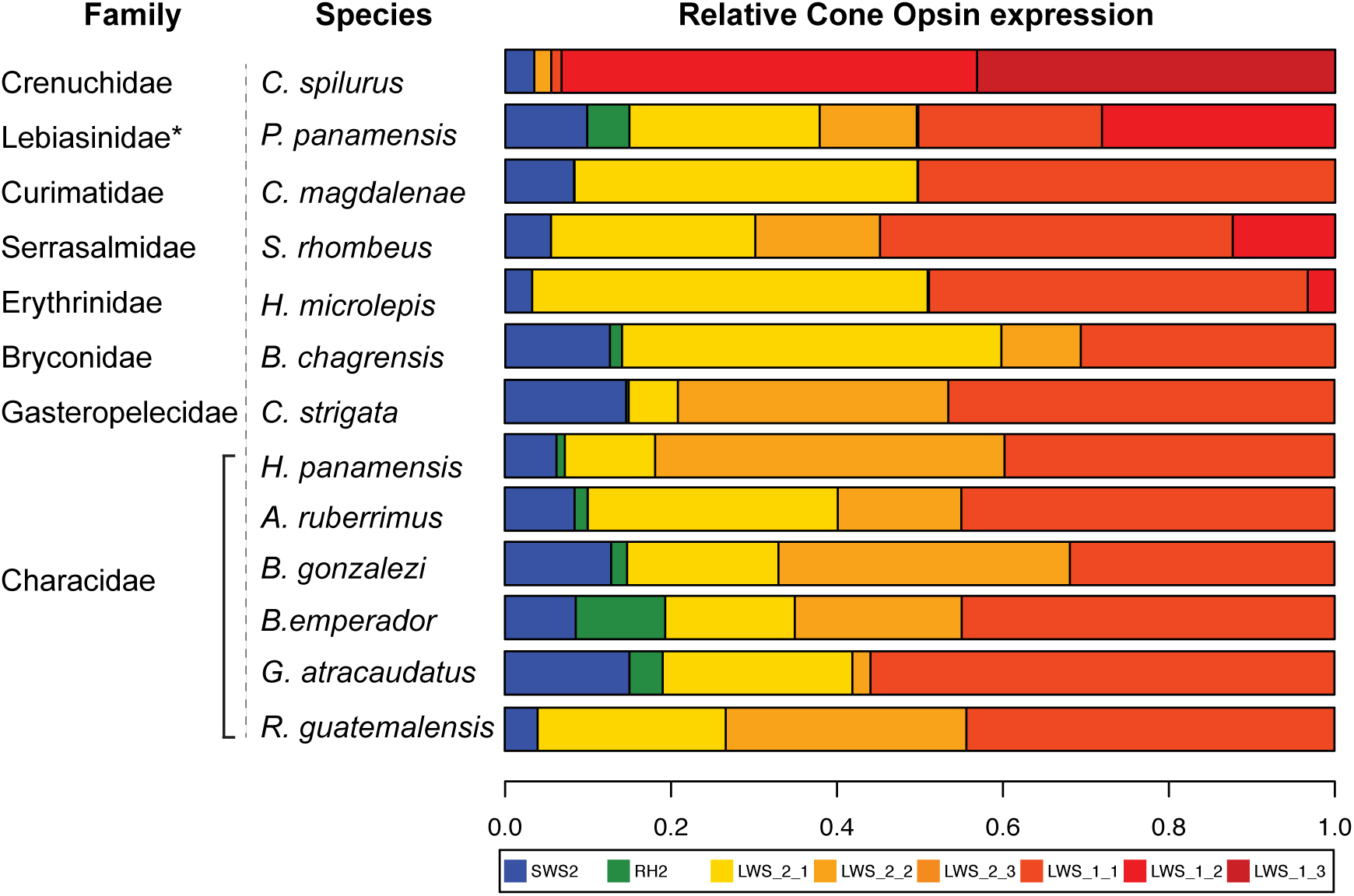
Opsin expression in Characiformes. Relative cone opsin expression is shown for each Characiform species and color-coded for each opsin.

The RH2 opsin was lowly expressed (<5%) in most samples, except in *B. emperador* (10%), and it was not recovered in the transcriptomes of four species (*C. spilurus, H. microlepis, S. rhombeus*, and *R. guatemalensis*). Finally, rod opsin expression was mainly dominated by the paralog RH1-1, while RH1-2 was very lowly expressed (<1.1%) in all analyzed species (Fig. S7).

### Photoreceptor spectral sensitivity

MSP of characiforms revealed a remarkable diversity in photoreceptor λ_max_. We identified up to six different cone classes based on spectral sensitivity: a blue-sensitive single cone (λ_max_=440-467 nm), a blue-green single cone (λ_max_=472-496 nm) and a second medium wavelength single cone, sensitive to the short-green (λ_max_=514-545 nm). Double cones contained either a green member (λ_max_=529-568 nm) paired with either a green-yellow (545-588 nm) or with a yellow-orange sensitive member (λ_max_=564-614 nm) (Fig. 7, Table 1). While all species showed at least three spectrally different photoreceptors, different species exhibited different sets. Rods exhibited similar variation within and across species, with a λ_max_ range of 502-536 nm.

**Table 1.** Cone and rod visual pigment peak sensitivities (λ_max_)

| Species | n | Photoreceptor type |  |  |  |  |  |  |
| --- | --- | --- | --- | --- | --- | --- | --- | --- |
|  |  | RH1 | SWS2 | RH2 | RH2/<br>LWS2 | LWS2 | LWS2/<br>LWS1 | LWS1 |
| Curimatidae |  |  |  |  |  |  |  |  |
| *C. magdalenae* | 3 | 517-536 | 450-455 | 476-496 | 535-545 | 531-554 | 588 | 585 |
| Erythrinidae |  |  |  |  |  |  |  |  |
| *H. microlepis* | 4 | 510-528 | — | 489-491 | 526-543 | 535-561 | 564-581 | 576-614 |
| Bryconidae |  |  |  |  |  |  |  |  |
| *B. chagrensis* | 4 | 504-531 | 446-467 | 472-485 | 515-535 | 532-568 | — | 564-612 |
| Characidae |  |  |  |  |  |  |  |  |
| *G. atracaudatus* | 3 | 504-516 | 440-459 | 491 | 521 | 530 | — | — |
| *B. gonzalezi* | 6 | 504-523 | 449-462 | 486-495 | 514-527 | 530-542 | 545 | 576 |
| *R. guatemalensis* | 6 | 502-423 | 447-466 | 481-495 | — | 530-541 | — | — |
| *A. ruberrimus* | 3 | 504-519 | 448-463 | 472-495 | 519-522 | 529-542 | — | — |

**Figure 7.**
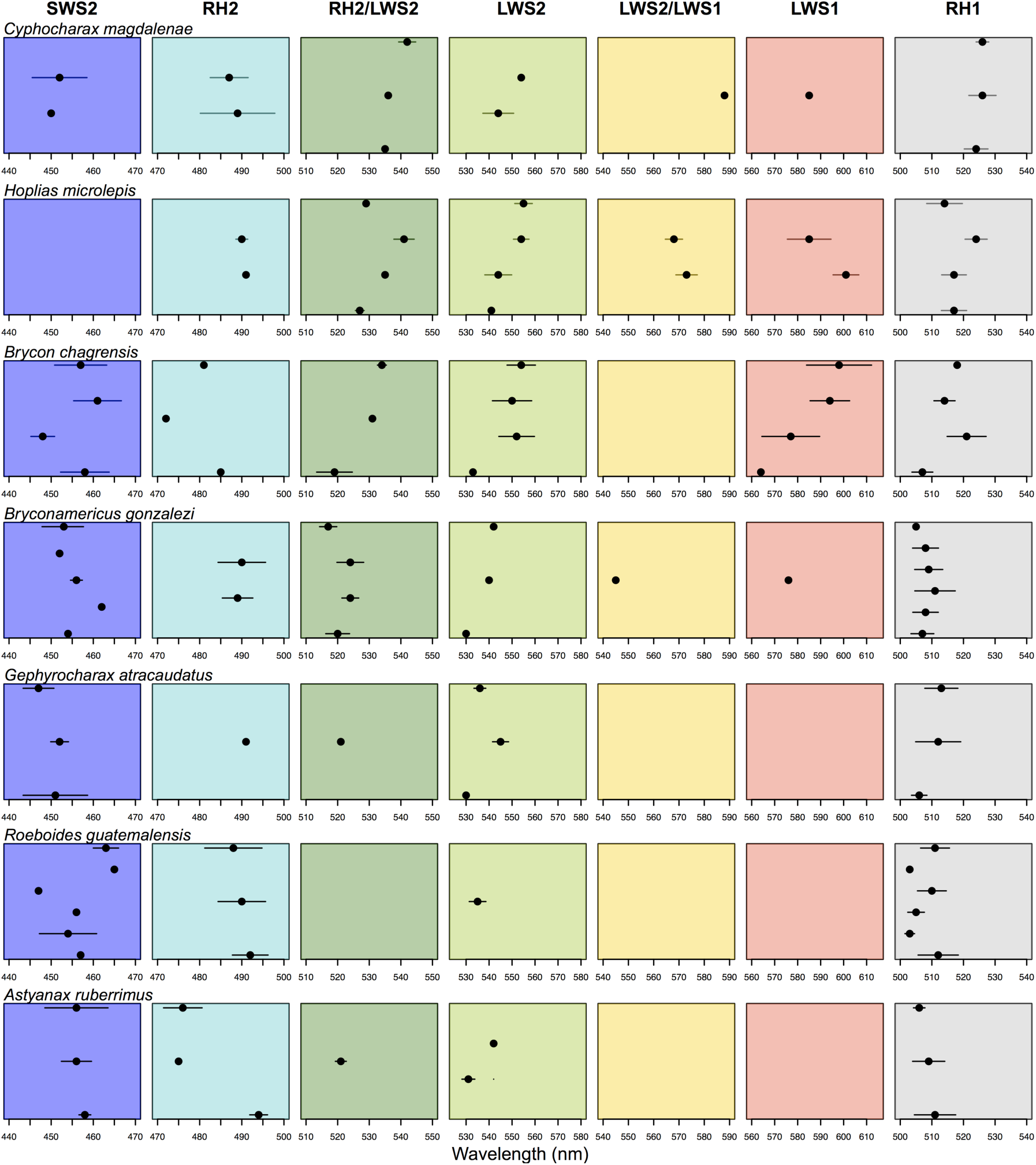
Microspectrophotometry of Characiformes. Black circles represent mean maximal absorbance (λ_max_) from visual pigments in wild-caught Panamanian characiforms. Horizontal lines denote mean standard deviation. Colored columns represent the six cone classes and rods. Each circle represents an individual analyzed for a given species.

Nomogram fitting and sequence analysis from this study, together with previous work on *Astyanax* (Parry et al. 2003) and other micro-spectrophotometric work on characiforms (Levine and Mac Nichol 1979), suggest that variation in λ_max_ in our dataset can be largely explained in terms of A_1_/A_2_ chromophore content (Fig. 8) with the exception of two groups that best fit a model including both coexpression and chromophore A_1_/A_2_ mixing. These two groups covered ranges of 514-545nm and 545-588nm and had shorter wavelength A_1_ sensitivities that were respectively best fit by a coexpressed RH2+LWS2 (in proportions 30%:70%) resulting in a 514nm predicted λ_maxA1_, and a coexpressed LWS2+LWS1 (in proportions 50%:50%) resulting in a 544nm predicted λ_maxA1_ (Table S4). The upper range for each of these groups was then fit by applying the same levels of vitamin A_2_ estimated for the other cone classes in the same individual to these coexpression λ_maxA1_ values. This provided models that consistently represented the best fit of the data, notably both across individuals and species. Records from these two classes could not be fit by any “pure” RH2 or LWS2, regardless of the amount of A_2_ imposed. It is important to observe that, these two classes were present in some but not all species, the coexpression λ_max_ values, assuming pure A_1_ (i.e. 514nm for RH2+LWS2; 544nm for LWS2+LWS1) allowed consistent fitting of similar uniform percentage values of vitamin A_2_ across classes within the same individual, and this held across individuals and species varying in their vitamin A_2_ content. Furthermore, based on the amino acid substitutions of the three key sites in the LWS1 opsin sequence (S164, Y261, T269), we predicted a λ_maxA1_ of 560 nm, which is in agreement with our MSP results. In addition, the LWS2 opsin exhibits substitutions (A164, F261, A269) with predicted λ_maxA1_ at 532 nm (Yokoyama et al. 2008) very close to the values obtained by MSP (529-530nm) from records of pure A_1_ photoreceptors (Table S4).

**Figure 8.**
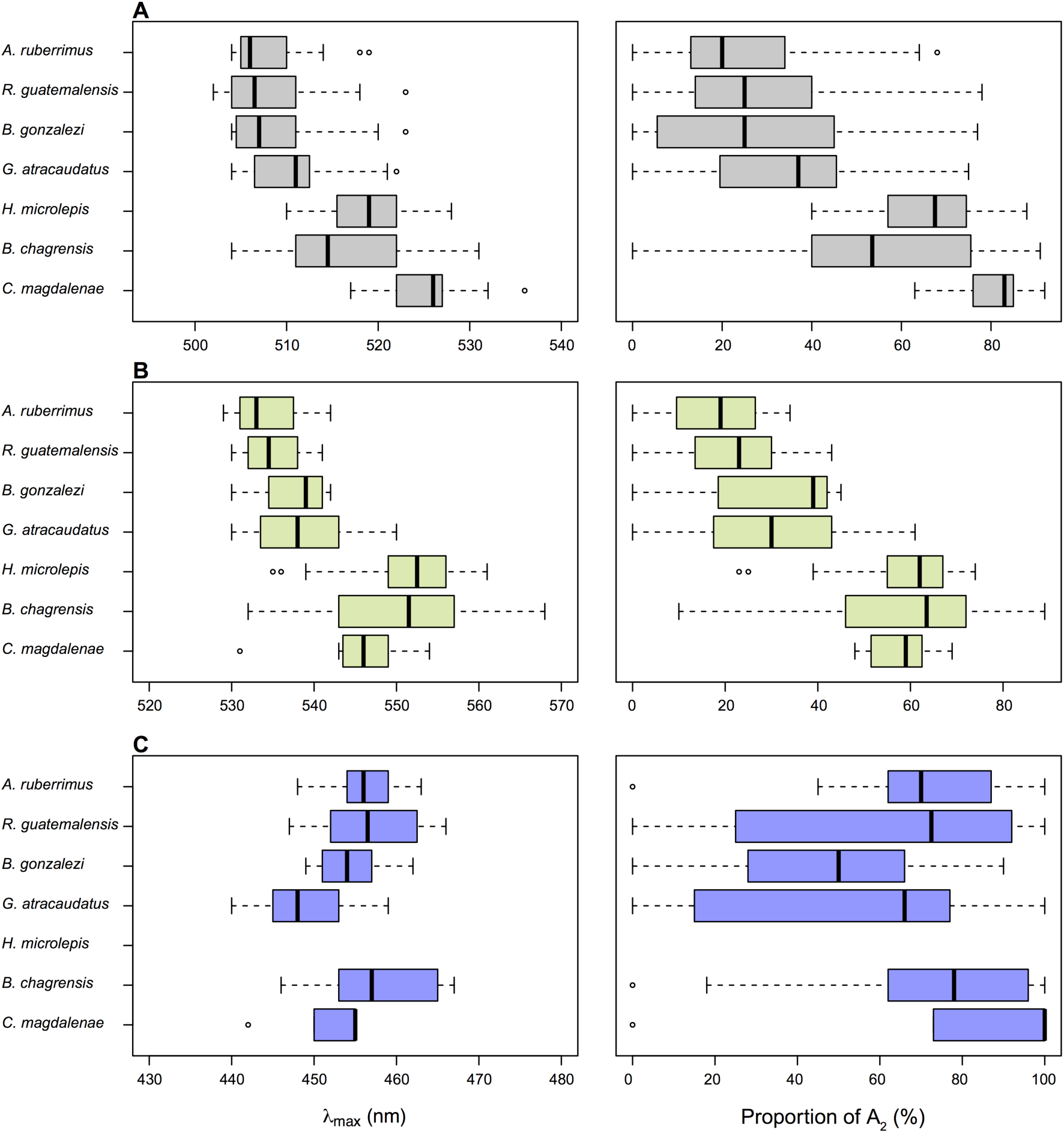
Boxplots showing variation in photoreceptor spectral sensitivity and proportion of vitamin A_2_ in Characiformes. (A) Rods. (B) LWS2 cones. (C) SWS2 cones.

### Genomic sequencing

Our phylogenetic tree based on genomic sequences of characiforms was consistent with results from previous studies (Oliveira et al. 2011), with African and Neotropical Characiformes sharing a monophyletic origin and being sister taxa to Gymnotiformes and Siluriformes (Fig. S9). All species used in this study were recovered to their expected taxonomic groups except for *Piabucina panamensis* which grouped with Parodontidae instead of Lebiasinidae.

## Discussion

### Dynamic opsin evolution in Characiformes

#### Opsin gene duplication and gene loss

Through transcriptome and genome analysis, we characterized opsin evolution in Neotropical Characiformes. Our results show that the opsin complement varies significantly between species (Fig. 3), with species utilizing from four (*R. guatemalensis*), to six cone opsins (*P. panamensis*). This diverse repertoire includes evidence for at least two separate copies of LWS opsins (LWS1 and LWS2), each with unique opsin sequences that originated after TGD. This was confirmed in our trees by the characiform LWS2 opsins clustering with the osteoglossimorph LWS2 opsins, which are known surviving copies of TGD (Liu et al. 2018). Our results also show that the LWS2 opsin underwent subsequent gene duplications within Characiformes, highlighting the susceptibility of opsin genes to duplication. LWS opsin gene duplications are not uncommon and they have independently occurred in several teleosts (Chinen et al. 2003; Matsumoto et al. 2006; Ward et al. 2008; Owens et al. 2009; Phillips et al. 2015; Liu et al. 2018). In addition, we also found another TGD surviving opsin product of a RH1 duplication: RH1-2. We confirmed this as the surviving duplicates clustered with the known RH1-2 opsins of Cypriniformes (Morrow et al. 2011; Morrow et al. 2017) (Fig. 2), however their functionality remains unknown. RH1-2 duplicates have also been found in the Japanese eel (*Anguilla japonica*) (Nakamura et al. 2017). Altogether, Characiformes have maintained TGD duplicates of two opsin classes (LWS and RH1) (Fig. S4), yet these duplicates have not been retained together in other teleosts.

The characiform visual pigment repertoire is also characterized by the absence of some opsin classes. The loss of SWS1 opsins seems to have occurred early in the evolution of Characiformes because its absence was shared between two phylogenetically distant species (*P. nattereri* and *A. mexicanus*). This is also corroborated by our gene expression and MSP data as we did not find any SWS1 cones (but see Parry et al. 2003). Additionally, RH2 seems to be variable across characiforms as it was absent in some species, but fully functional in others; a result supported by both MSP and gene expression.

#### Opsin gene conversion

We found evidence of gene conversion in both LWS1 and LWS2 opsins. GARD analysis showed that the recombination locations are primarily in the first exons, which suggests there might be selective pressures preventing gene conversion from homogenizing coding sequences where the “key sites” are located in exons 3, 4 and 5. Gene conversion is further supported by the different tree-topologies, a pattern more evident in LWS2 where the tree based on exons 3 to 6 (Fig. S5B) suggests LWS2 opsins duplicated in the early ancestor of Bryconidae, Gasteropelecidae and Characidae. This is a more parsimonious pattern than LWS2 duplications occurring in each family independently and is also in agreement with our tree based on genomic sequencing (Fig. S9). Overall, our findings are supported by other studies reporting gene conversion homogenizing opsin sequences (Watson et al. 2010; Rennison et al. 2012; Nakamura et al. 2013; Cortesi et al. 2015; Escobar-Camacho et al. 2017). However, it has also been suggested that gene conversion might increase allelic diversity (Ohta 2010).

### Opsin neofunctionalization: evolution of spectral tuning and opsin gene expression

The great diversity in visual sensitivities among Characiformes is the product of different spectral tuning mechanisms acting in concert. These include opsin sequence tuning, opsin gene expression, opsin gene loss and duplication, opsin coexpression, and chromophore tuning.

#### Opsin sequence tuning evolution

As discussed above, some characiforms have lost the RH2 opsin through opsin downregulation and pseudogenization. However, characiforms have expanded their green sensitivity by utilizing another opsin class gained from opsin gene duplication followed by opsin sequence tuning. Through genetic and electrophysiology experiments we confirm that LWS2 opsins are sensitive to green light and that this is maintained in all analyzed species (Fig. 5,7). This is consistent with previous molecular studies that showed that *Astyanax* had green sensitive opsins (Yokoyama and Yokoyama 1990b; Yokoyama and Yokoyama 1990a) due to mutations in three of the known “five-sites” (Yokoyama and Radlwimmer 1998). In our analysis, the diversity at spectral tuning sites in the LWS opsins, particularly site 164 (Fig. S6), showed the ability of opsins to acquire new functions through opsin sequence variation. This is important because shifts in λ_max_ can have profound impacts on fish color vision. As the λ_max_ of a photoreceptor shifts across the wavelength spectrum, chromatic contrast will also vary in the visual color space and this could affect chromatic discrimination.

In addition, micro-spectrophotometry was not able to distinguish the two LWS2 (LWS2-1 and LWS2-2) duplicates and it remains unclear whether they differ at all in λ_max_. Protein reconstitution might shed light on the effects of the observed substitutions in the two LWS2 opsins and whether these have an influence on spectral absorbance.

The presence of LWS2 opsins in both Osteoglossiformes and Characiformes suggests that LWS2-green sensitivity evolved in an early ancestor dated after TGD but before the split of Osteoglossomorpha and Clupeocephala (around 240 MYA) (Hughes et al. 2018). This implies that LWS2 opsins have been maintained in Osteoglossiformes and Characiformes for at least over 300 million years (Liu et al. 2018), while they have been lost in several other teleosts. Indeed, there is evidence that most duplicated genes were lost in the first 60 million years after TGD (Inoue et al. 2015). Green sensitivity might have evolved during the Permian (∼300-250 MYA), a period characterized by several fish extinction events in its early and middle epochs and ending with the Permian mass extinction around 251 MYA (Romano et al. 2016). Therefore, the LWS2 neofunctionalization through opsin sequence tuning might be a result of the strong environmental changes characterizing the Permian where green sensitivity was favored in collapsed freshwater environments.

#### Opsin expression

Novel opsins can also acquire new functions through gene expression mechanisms (Cortesi et al. 2015). In Characiformes, it seems that the rise of the LWS2 opsins might have changed the regulatory architecture of RH2 expression, leading to downregulation and even to gene loss (Fig. 6,S4). Previous studies have found the same pattern between two different opsin classes: whenever a strong shift in λ_max_ occurs in one opsin, there can be gene loss/downregulation in another one. In flounder and South American cichlids, the SWS2 opsin has acquired green sensitivity, while the functionality of the RH2 opsins has been reduced (Kasagi et al. 2018, Escobar-Camacho et al. 2019). A similar pattern has occurred in Osteoglossiformes where the LWS2 opsin is green sensitive and the RH2 opsin has been lost (Liu et al. 2018).

Micro-spectrophotometry suggests that, in double cones, the LWS2 opsin is sometimes coexpressed in RH2/LWS2 mixes or in LWS2/LWS1. Immunohistochemistry and/or in-situ approaches will be needed to characterize in more detail the extent and localization in the retina of such coexpressed opsins. Coexpression of RH2 and LWS opsins has been observed in mammals (Applebury et al. 2000; Parry and Bowmaker 2002; Lukáts et al. 2005), amphibians (Isayama et al. 2014), in the guppy (Archer and Lythgoe 1990) and appears to be widespread in another highly diverse freshwater group; the African and Neotropical cichlids (Dalton et al. 2014; Dalton et al. 2015; Torres-Dowdall et al. 2017).

Regional variation in the expression of RH2, single or coexpressed, is likely at the origin of the discrepancy in its abundance as measured by whole-retina transcriptomics and by micro-spectrophotometry. MSP is a useful approach to identify the peak of spectral sensitivity of a photoreceptor, particularly when coexpression and/or chromophore variation are present, i.e. when opsin sequence does not provide sufficient information to infer photoreceptor sensitivity. However, MSP samples a limited number of photoreceptors in the regions of the retina that happen to be scanned, so it is not appropriate for quantitative estimates of photoreceptor or opsin class abundances.

Our results show that there is differential opsin expression in Characiformes. In Characidae, most species express LWS2-2 more than LWS2-1, whereas this is the opposite in species from other families (Fig. 6). Even though our data set is based on a few individuals and more sampling is needed to quantify differential opsin expression, this suggests there might be a pattern in which opsins are differentially regulated in different species. In addition, we also found significant differential expression between RH1-1 and RH1-2 (Fig. S7) yet we do not know whether RH1-2 has a specific role in the visual system. More studies are needed to fully characterize its functionality. It has been shown that RH1-2 duplicates can acquire specific functions such as regionalized expression in the zebrafish retina (Morrow et al. 2017), or ontogenetic expression in the life cycle of the Japanese eel (Nakamura et al. 2017).

#### Chromophore tuning

Fish visual pigments can be based on alternative chromophores, either 11-cis retinal, a derivative of vitamin A_1_, or 3,4-didehydroretinal, a derivative of vitamin A_2,_ or on mixtures of the two, with the potential to generate large shifts in photoreceptors λ_max_ at longer wavelengths (Parry and Bowmaker 2000). In our dataset the spectral sensitivities of rods and cones are significantly different between species sampled in murky waters (*B. chagrensis, C. magdalenae* and *H. microlepis*) vs. clear-waters (*A. ruberrimus, B. gonzalezi, G. atracaudatus*, and *R. guatemalensis*) (Fig. 8, Table S5-6), with the red-shift attributable mainly to high levels of vitamin A_2_ in species sampled in turbid environments (Fig. 8). Since variation in spectral sensitivities is due, to a large extent, to differences in vitamin A_2_ content, it is not surprising that, with the notable exception of *G. atracaudatus*, there was little variation in blue cone sensitivity across species, as the effects of the chromophore are minimal at short wavelengths (Whitmore and Bowmaker 1989). Chromophore-based spectral tuning is characteristic of fish inhabiting long-wavelength-shifted habitats (Whitmore and Bowmaker 1989; Carleton et al. 2006; Toyama et al. 2008; Hofmann et al. 2009; Miyagi et al. 2012; Saarinen et al. 2012; Weadick et al. 2012; Liu et al. 2016; Terai et al. 2017; Torres-Dowdall et al. 2017; Escobar-Camacho et al. 2019), and is well suited to the variable light environments of Neotropical freshwater ecosystems, famously among the most diverse in spectra on the planet, from clear fast-running mountain creeks, to muddy ‘white waters’ (from ‘*agua blanca’*, indicating turbid highly scattering brown-tainted waters), and tannin-rich black waters (Wallace 1865; Costa et al. 2013; Escobar-Camacho et al. 2019). Finally, non-significant variation in rod chromophore proportions was observed between individuals, with some species exhibiting a similar proportion across individuals, while other species exhibit a larger range (Table S7). Variable A_1_/A_2_ ratios have been previously found in the characid *A. fasciatus* (Parry et al. 2003), and in rods of several other characiforms (Schwanzara 1967; Levine & MacNichol 1979).

### Opsins and characiform phylogenetics

Even though our genomic multilocus phylogeny suggested a monophyletic origin of Characiformes, including African and Neotropical lineages (Citharinoidei and Characoidei respectively) (Fig. S9), several of our opsin trees contradict this pattern because Characiformes appear paraphyletic in relation to Siluriformes and Gymnotiformes (Fig. S1-3). These contrasting results are not surprising as the non-monophyly of Characiformes has been reported before (Nakatani et al. 2011; Chen et al. 2013; Hakrabarty et al. 2017), although; other comprehensive studies have resolved Characiformes as monophyletic (Betancur-R et al. 2013; Arcila et al. 2017; Hughes et al. 2018). Studies that find Characiformes paraphyletic often find discrepancies between Citharinoidei and Characoidei, where the latter often clusters as sister group to Siluriformes (Nakatani et al. 2011; Chen et al. 2013; Hakrabarty et al. 2017). Interestingly, we obtained paraphyletic results in our opsin trees because opsin sequences from Gymnotiformes and Siluriformes clustered within the opsin clades of Characiformes, although we did not include opsin sequences of African species. The opsin tree topologies of this study could be the result of substitution saturation over evolutionary time or indeed a signal of paraphyletic origins of Characiformes. Future studies should include analysis of opsins of African characiforms in order to elucidate this pattern.

## Conclusions

Through molecular and electrophysiological experiments we have characterized the visual system of Neotropical Characiformes. Their opsin repertoire is a product of complex evolutionary dynamics characterized by opsin gene loss (SWS1, RH2) and opsin gene duplication (LWS and RH1). These opsin duplicates are a product of a teleost whole genome duplication (TGD) and from characiform-specific duplication events that have been maintained for hundreds of millions of years. The LWS duplicates have acquired new functions through amino acid substitution in key sites that shift their maximal absorbance to green light. These duplicates exhibit gene conversion, and utilize variable codons in key tuning sites leading to reversion and parallel evolution. In addition, the SWS2 opsin exhibits great amino acid variation across species that might shift spectral sensitivities, and the RH1-2 opsin has a different pattern in opsin expression as it is always downregulated in our samples.

The diversity of visual pigments in Characiformes is the product of several spectral tuning mechanisms acting in concert. These are mainly opsin sequence variation, opsin gene loss and duplication, and A_1_/A_2_ chromophore tuning. Such mechanisms have probably allowed characiforms to thrive in the variable freshwater light environments of the Neotropics. Overall, the visual system of Characiformes showcases how opsins acquire new functions and the divergent evolutionary pathway followed by this group compared to other teleosts. This study shows how studying speciose, understudied groups, provides a unique opportunity to better understand opsin gene evolution and spectral tuning mechanisms.

## Materials and methods

### Animals

Adult fish specimens were collected using fishing lines, manual seines and cast nets in several locations in Panama and Suriname from May to July of 2017. Fish were caught either in murky, black, or clear waters (Table S8). Sampling permits were in accordance with the Panamanian and Suriname laws of environmental protection (permits from Ministerio de Ambiente de Panamá, MiAmbiente, permit No. SC/A-14-17; and Ministry of Agriculture, Animal Husbandry and Fisheries of Suriname, permit No. 1087). Fish were handled following STRI IACUC protocol (#2017-0501-2020). After sampling all fish were brought back to the Naos Research Laboratories at the Smithsonian Research Institute Panama. Three specimens from each species were killed immediately for RNA-Sequencing and a total of 29 specimens were used for micro-spectrophotometry (MSP). Fish sampled in Suriname were sacrificed at the laboratories of Anton de Kom University of Suriname. In total, we obtained 13 species that belonged to eight different families within Characiformes (Tables S8).

### RNA seq

After collection, fish were euthanized with buffered MS-222, their eyes were enucleated and their retinas preserved in RNAlater. Two or three samples per species were used for RNA sequencing. Total RNA was extracted with an RNeasy kit (Qiagen) and RNA quality was verified on an Agilent Bioanalyzer. RNAseq libraries were prepared using the Illumina TruSeq RNA library preparation kit (Illumina Inc, San Diego) and sequenced to obtain 100-bp paired-end-reads with a total of 36 samples multiplexed in three lanes (12 samples per lane) on an Illumina HiSeq1500 sequencer at the University of Maryland Institute for Bioscience & Biotechnology Research. The quality of the data was checked using FastQC version 0.11.2. Further, we used Trimmomatic version 0.32 (Bolger et al. 2014) to remove overrepresented sequences and to retain sequences with a minimum quality score of 20 and a minimum length of 80 bp. Transcriptomes were combined to obtain 13 de-novo assemblies for each species. This was performed with Trinity version r20140413 (Haas et al. 2013), using only paired sequences with a minimum coverage of two to join contigs.

### Opsin phylogenetics and molecular analysis

Candidate opsin sequences were identified from the assembled transcriptome FASTA files by Tblastx querying with the opsin genes of *Astyanax fasciatus* (Yokoyama and Yokoyama 1990a; Yokoyama and Yokoyama 1993; Register et al. 1994; Yokoyama et al. 1995). Because we found opsin duplicates in the transcriptomes, we used GENEIOUS 8.1 to map paired-reads for each paralog and correctly assemble each opsin sequence. We confirmed the identities of gene sequences for each species to a particular opsin class based on their phylogenetic relationships with opsins sequences of lamprey (*Geotria australis*) and several teleosts obtained through Genbank (Benson et al. 2005). Furthermore, we added to our analysis opsin sequences of the available genomes of the Mexican cavefish, (*Astyanax mexicanus*), and the red-bellied piranha (*Pygocentrus nattereri*). We used MAFFT (Katoh et al. 2002) to align amino acid sequences and ProtTest 3.4.2 (Darriba et al. 2011) to obtain the evolutionary models for each opsin class (Table S9). We used RAXML for building maximum-likelihood trees. We ran 10 searches for the best tree and performed 1000 bootstrap replicates in RAXML 8.0 on CIPRES (Miller et al. 2015).

Once we identified the characiform opsin classes, we searched for amino acid subtitutions that could shift the spectral sensitivity of visual pigments. To do this, we aligned characiform opsin sequences with bovine rhodopsin and with opsins from other teleosts. We looked for substitutions that fell in putative transmembrane regions and in the retinal binding pocket facing the chromophore or in known spectral tuning sites (Hunt et al. 2001; Carleton et al. 2005; Yokoyama 2008; Yokoyama et al. 2008).

Finally, since there were opsin duplicates in most analyzed species, we tested whether there was gene conversion, as this has recently emerged as a common phenomenon in teleosts (Owens et al. 2009; Watson et al. 2010; Nakamura et al. 2013; Escobar-Camacho et al. 2017; Sandkam et al. 2017). For this we used the program GARD (Genetic Algorithm Recombination Detection) (Kosakovsky Pond et al. 2006) on separate alignments of the LWS duplicates to detect the presence or absence of recombination. To corroborate patterns of gene conversion we built phylogenetic trees based on the fragments between the recombination breakpoints.

### Ancestral state reconstruction

Previous research has characterized the molecular basis of spectral tuning in the LWS pigments where five amino acid changes (S164A, H181Y, Y261F, T269A, A292S) can shift λ_max_ up to 50 nm (Yokoyama and Yokoyama 1990a; Yokoyama and Radlwimmer 1998; Yokoyama and Radlwimmer 2001; Yokoyama et al. 2008). Since we observed variation in the occurrence of three of these amino-acid substitutions (S164A, Y261F, T269A) that are known to shift to short wavelengths the λ_max_ of LWS opsins by 7, 10 and 16 nm respectively (Asenjo et al. 1994; Takahashi and Ebrey 2003; Yokoyama 2008; Yokoyama et al. 2008), we analyzed the evolutionary relationships between the spectral tuning sites across the Characiformes. We observed seven combinations of the three sites in our LWS dataset (Table S10). We assigned one of these combinations to each LWS opsin gene and performed a discrete trait ancestral state reconstruction analysis in R (R Core Team 2014).

In addition, we observed that, among the three tuning sites, site S164A was the most variable. In order to understand the molecular mechanisms leading to this variation, we reconstructed the evolutionary changes leading to both serine or alanine and identified parallel changes and reversions by characterizing the extant codons in each gene and performing ancestral state reconstruction where we incorporated the respective codon to each gene as a trait.

For ancestral state reconstruction analyses we used the ace function in the APE package (Paradis and Schliep 2018) in R. The ace function employs a maximum likelihood approach where the reconstructed ancestral states are given as a proportion of the total likelihood for each state for each node.

### Opsin gene expression

For estimating gene expression of each opsin, reads were mapped back to the assembled transcriptomes using RSEM as part of the Trinity package (Haas et al. 2013). Read counts for each opsin class were extracted from RSEM output (quantified as fragments per kilobase of transcript per million reads, FPKM). In order to avoid non-independent bias of opsin expression owing to variation in the expression of each opsin class, cone opsin read counts were then normalized to those of the β-actin gene. We also normalized for total cone opsin expression and divided the expression of each opsin by the sum of all cone opsin counts to get the proportion of each expressed opsin.

### Photoreceptor spectral sensitivity

The peak of maximum spectral sensitivity of individual photoreceptors was obtained by micro-spectrophotometry of fresh retinas from wild-caught fish from the same collecting sites as the individuals used for gene expression analysis. Fish were dark-adapted for at least 2 hrs, after which they were sacrificed with an overdose of buffered MS222. Eyes were enucleated under a dissecting scope in dim deep red light. The retina was removed and transferred to a PBS solution containing 6.0% sucrose (Sigma). A small piece of retina was cut out and placed on a glass cover slip in a drop of solution, then delicately macerated with razor blades. The preparation was covered with a second glass cover slip and sealed with high-vacuum silicone grease (Dow Corning) (Escobar-Camacho et al. 2019).

Spectral absorbance was measured with a computer-controlled single-beam micro-spectrophotometer fitted with quartz optics and a 100W quartz-halogen lamp (Loew 1982). Baseline records were taken by averaging a scan from 750 to 350 nm and a second in the opposite direction, through a clear area of the preparation and in proximity to the photoreceptor of interest. A record of the visual cell was then obtained by scanning with the MSP beam through the photoreceptor outer segment. Finally, the cell’s absorption spectrum was obtained by subtracting the baseline record. A custom-designed spectral analysis program (Loew et al. unpublished data) was used to determine λ_max_ from absorbance records using existing templates (Dartnall 1953; Munz and Schwanzara 1967). Individual spectra were smoothed with a nine-point adjacent averaging function and the resulting curves were differentiated to obtain a preliminary maximum value. This was used to normalize curves to zero at the baseline on the long wavelength limb and to one at the maximum value (Escobar-Camacho et al. 2019). Whitmore and Bowmaker’s (1989) relationship (Eqn 1) was used to recursively fit the observed (normalized) absorption spectra to curves resulting from combinations of different proportions of pure Vitamin A_1_ and corresponding pure Vitamin A_2_ nomograms (Whitmore and Bowmaker 1989).

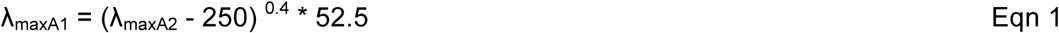

A non-parametric ANOVA on ranks (Kruskal-Wallis t test) followed by a pairwise Wilcoxon test was used to compare the λ_max_ of different photoreceptor classes.

### DNA extraction, sequencing and phylogenetic analysis

To analyze the evolutionary relationships of the sampled characiforms and confirm species identification, we sequenced nuclear and mitochondrial genes (16S, Cytb, Myh6, RAG1 and RAG2) from all collected species except *C. spilurus*. DNA was extracted with a DNeasy kit (Qiagen), and DNA quality was verified using the Nanodrop approach. PCR reactions were performed in a total volume of 50µl using a thermocycler (Eppendorf). Reactions contained 25µl DreamTaq DNA polymerase, 20µl of sterile distilled H_2_O, 2µl of each primer (10µM), and 1µl template DNA. Conditions were as follow: 94°C (2 min); 35 cycles of 94°C (30s), °54 (30s), and 72°C (2 min) followed by 72°C (4min). Nested-PCRs were used to amplify the genes RAG1 and RAG2. Amplified products were checked on 1% agarose gel stained with GelRed.

Once individual genes were sequenced for each species, all genes were concatenated. We added our species alignments to a data set of 213 sequenced characiforms for the same markers from Oliveira et al. (2011). We used Partitionfinder2 (Lanfear et al. 2017) to obtain the most appropriate phylogenetic models and partitioning scheme. Finally, we used RAXML to build maximum-likelihood trees with 1000 Bootstrap repetitions (Miller et al. 2015).

## Acknowledgements

Special thanks go to Suwei Zhao for training during library preparations. We thank the University of Maryland Institute for Bioscience & Biotechnology Research for sequencing. We thank Michaela Taylor for help during genomic sequencing and Danielle Adams for help during ancestral reconstruction analysis. We are grateful to Ellis R. Loew for generously providing his MSP machine and analyses software. We thank Alejandra Rodríguez-Abaunza and Aureliano Valencia for their assistance during sampling. We also thank all of the staff at Bocas del Toro Research Station and at Naos Laboratories, Smithsonian Tropical Research Institute (STRI), Panama, for their help during our field season. We also thank Owen McMillan for logistic support and Richard Cooke for his valuable insight during our field season. This work was supported by a STRI Short Term Fellowship (ID 102755 to D.E-C); the National Institute of Health (R01EY024693 to K.L.C) and by the Secretariat of Higher Education, Science, and Technology and Innovation of Ecuador (SENESCYT) (2014-AR2Q4465 to D.E-C).

## Data availability

DNA sequences and transcriptome libraries will be available upon manuscript acceptance.

