## Supplementary materials for "Visual pigment evolution in Characiformes: the dynamic interplay of teleost whole-genome duplication, surviving opsins and spectral tuning"

### Supplementary Tables

**Table S1.** Recombination points obtained from GARD

| | | |
|---|---|---|
| **LWS1** | | |
| Breakpoint 1 | Breakpoint 2 | |
| 1- 251 bp | 252-1074 bp | |
| **LWS2** | | |
| Breakpoint 1 | Breakpoint 2 | Breakpoint 3 |
| 1-209 bp | 210-404 bp | 405-1065 bp |

**Table S2.** Likelihoods of ancestral state reconstruction for LWS2 amino-acid combinations

| **Nodes** | **Combinations** | | | | | | |
|---|---|---|---|---|---|---|---|
| | 1(Blue) | 2(Yellow) | 3(Purple) | 4(Pink) | 5(Cyan) | 6(Green) | 7(Magenta) |
| **1** | 94.155 | 2.219 | 0.731 | 0.671 | 0.881 | 0.671 | 0.671 |
| **2** | 73.564 | 13.230 | 2.686 | 2.259 | 3.743 | 2.259 | 2.259 |
| **3** | 97.450 | 0.087 | 0.087 | 0.087 | 2.113 | 0.088 | 0.087 |
| **4** | 99.700 | 0.047 | 0.047 | 0.047 | 0.060 | 0.051 | 0.047 |
| **5** | 98.735 | 0.056 | 0.056 | 0.056 | 0.789 | 0.253 | 0.056 |
| **6** | 99.746 | 0.042 | 0.042 | 0.042 | 0.042 | 0.042 | 0.042 |
| **7** | 99.809 | 0.032 | 0.032 | 0.032 | 0.032 | 0.032 | 0.032 |
| **8** | 99.950 | 0.008 | 0.008 | 0.008 | 0.008 | 0.008 | 0.008 |
| **9** | 99.746 | 0.042 | 0.042 | 0.042 | 0.042 | 0.044 | 0.042 |
| **10** | 99.489 | 0.070 | 0.070 | 0.070 | 0.075 | 0.154 | 0.070 |
| **11** | 96.510 | 0.542 | 0.542 | 0.542 | 0.779 | 0.542 | 0.542 |
| **12** | 97.775 | 0.070 | 0.070 | 0.070 | 1.877 | 0.070 | 0.070 |
| **13** | 44.620 | 0.986 | 0.986 | 0.986 | 50.451 | 0.986 | 0.986 |
| **14** | 0.060 | 0.060 | 0.060 | 0.060 | 99.637 | 0.060 | 0.060 |
| **15** | 95.896 | 0.288 | 0.288 | 0.288 | 0.288 | 2.662 | 0.288 |
| **16** | 60.843 | 0.966 | 0.966 | 0.966 | 0.966 | 34.327 | 0.966 |
| **17** | 99.927 | 0.012 | 0.012 | 0.012 | 0.012 | 0.012 | 0.012 |
| **18** | 99.999 | 0.000 | 0.000 | 0.000 | 0.000 | 0.000 | 0.000 |
| **19** | 99.945 | 0.009 | 0.009 | 0.009 | 0.009 | 0.009 | 0.009 |
| **20** | 99.938 | 0.010 | 0.010 | 0.010 | 0.010 | 0.010 | 0.010 |
| **21** | 99.953 | 0.006 | 0.006 | 0.006 | 0.016 | 0.006 | 0.006 |
| **22** | 99.631 | 0.062 | 0.062 | 0.062 | 0.062 | 0.062 | 0.062 |
| **23** | 99.316 | 0.036 | 0.036 | 0.036 | 0.505 | 0.036 | 0.036 |
| **24** | 81.896 | 0.489 | 0.489 | 0.489 | 15.658 | 0.489 | 0.489 |
| **25** | 99.994 | 0.001 | 0.001 | 0.001 | 0.001 | 0.001 | 0.001 |
| **26** | 100.000 | 0.000 | 0.000 | 0.000 | 0.000 | 0.000 | 0.000 |
| **27** | 99.917 | 0.014 | 0.014 | 0.014 | 0.014 | 0.014 | 0.014 |
| **28** | 100.000 | 0.000 | 0.000 | 0.000 | 0.000 | 0.000 | 0.000 |
| **29** | 99.992 | 0.001 | 0.001 | 0.001 | 0.001 | 0.001 | 0.001 |
| **30** | 99.778 | 0.037 | 0.037 | 0.037 | 0.037 | 0.037 | 0.037 |
| **31** | 99.754 | 0.041 | 0.041 | 0.041 | 0.041 | 0.041 | 0.041 |
| **32** | 58.954 | 1.214 | 1.214 | 1.214 | 27.831 | 8.360 | 1.214 |
| **33** | 1.541 | 1.541 | 1.541 | 1.541 | 71.870 | 20.424 | 1.541 |
| **34** | 99.459 | 0.090 | 0.090 | 0.090 | 0.090 | 0.090 | 0.090 |
| **35** | 96.753 | 0.541 | 0.541 | 0.541 | 0.541 | 0.541 | 0.541 |
| **36** | 99.944 | 0.009 | 0.009 | 0.009 | 0.009 | 0.009 | 0.009 |
| **37** | 35.894 | 1.410 | 1.410 | 1.410 | 57.056 | 1.410 | 1.410 |
| **38** | 99.898 | 0.017 | 0.017 | 0.017 | 0.017 | 0.017 | 0.017 |
| **39** | 99.052 | 0.158 | 0.158 | 0.158 | 0.158 | 0.158 | 0.158 |
| **40** | 0.285 | 0.285 | 0.285 | 0.285 | 98.288 | 0.285 | 0.285 |
| **41** | 0.272 | 0.272 | 0.272 | 0.272 | 98.368 | 0.272 | 0.272 |
| **42** | 0.003 | 0.000 | 0.000 | 0.000 | 99.997 | 0.000 | 0.000 |
| **43** | 0.438 | 93.743 | 4.065 | 0.439 | 0.438 | 0.438 | 0.438 |
| **44** | 0.002 | 99.985 | 0.002 | 0.007 | 0.002 | 0.002 | 0.002 |
| **45** | 0.062 | 97.261 | 0.062 | 2.428 | 0.062 | 0.062 | 0.062 |
| **46** | 0.673 | 58.987 | 0.673 | 37.647 | 0.673 | 0.673 | 0.673 |
| **47** | 0.073 | 99.559 | 0.073 | 0.073 | 0.073 | 0.073 | 0.073 |
| **48** | 0.000 | 100.000 | 0.000 | 0.000 | 0.000 | 0.000 | 0.000 |
| **49** | 0.000 | 100.000 | 0.000 | 0.000 | 0.000 | 0.000 | 0.000 |
| **50** | 0.018 | 99.140 | 0.111 | 0.567 | 0.018 | 0.018 | 0.128 |
| **51** | 0.538 | 55.862 | 5.574 | 30.435 | 0.538 | 0.538 | 6.515 |
| **52** | 0.872 | 0.872 | 13.671 | 66.775 | 0.872 | 0.872 | 16.065 |
| **53** | 0.216 | 0.216 | 16.233 | 82.689 | 0.216 | 0.216 | 0.216 |
| **54** | 0.000 | 0.000 | 100.000 | 0.000 | 0.000 | 0.000 | 0.000 |
| **55** | 0.022 | 0.022 | 99.867 | 0.022 | 0.022 | 0.022 | 0.022 |
| **56** | 0.006 | 0.006 | 0.006 | 99.963 | 0.006 | 0.006 | 0.006 |
| **57** | 0.010 | 99.796 | 0.010 | 0.155 | 0.010 | 0.010 | 0.010 |

| | | | | | | | |
|---|---|---|---|---|---|---|---|
| **58** | 0.196 | 90.904 | 0.196 | 8.117 | 0.196 | 0.196 | 0.196 |
| **59** | 0.004 | 99.978 | 0.004 | 0.004 | 0.004 | 0.004 | 0.004 |
| **60** | 0.000 | 100.000 | 0.000 | 0.000 | 0.000 | 0.000 | 0.000 |
| **61** | 0.002 | 99.985 | 0.002 | 0.002 | 0.002 | 0.002 | 0.002 |
| **62** | 0.000 | 100.000 | 0.000 | 0.000 | 0.000 | 0.000 | 0.000 |
| **63** | 0.038 | 98.958 | 0.038 | 0.852 | 0.038 | 0.038 | 0.038 |
| **64** | 1.588 | 40.224 | 1.588 | 51.834 | 1.588 | 1.588 | 1.588 |
| **65** | 0.035 | 99.792 | 0.035 | 0.035 | 0.035 | 0.035 | 0.035 |
| **66** | 0.007 | 99.960 | 0.007 | 0.007 | 0.007 | 0.007 | 0.007 |
| **67** | 0.004 | 99.974 | 0.004 | 0.004 | 0.004 | 0.004 | 0.004 |
| **68** | 0.023 | 99.862 | 0.023 | 0.023 | 0.023 | 0.023 | 0.023 |
| **69** | 0.005 | 99.970 | 0.005 | 0.005 | 0.005 | 0.005 | 0.005 |
| **70** | 0.007 | 99.956 | 0.007 | 0.007 | 0.007 | 0.007 | 0.007 |
| **71** | 0.042 | 99.746 | 0.042 | 0.042 | 0.042 | 0.042 | 0.042 |
| **72** | 0.005 | 99.971 | 0.005 | 0.005 | 0.005 | 0.005 | 0.005 |
| **73** | 0.000 | 100.000 | 0.000 | 0.000 | 0.000 | 0.000 | 0.000 |
| **74** | 0.007 | 99.957 | 0.007 | 0.007 | 0.007 | 0.007 | 0.007 |
| **75** | 0.000 | 100.000 | 0.000 | 0.000 | 0.000 | 0.000 | 0.000 |
| **76** | 2.276 | 50.863 | 37.758 | 2.276 | 2.276 | 2.276 | 2.276 |
| **77** | 0.317 | 98.100 | 0.317 | 0.317 | 0.317 | 0.317 | 0.317 |

**Table S3.** Likelihoods of ancestral state reconstruction for the amino-acid site S164A

| Node | Codon | | | | | |
|---|---|---|---|---|---|---|
| | 1 (GCT) | 2 (TCT) | 3 (TCC) | 4 (GCA) | 5 (TCA) | 6 (GCC) |
| **1** | 2.11262 | 83.90329 | 4.24989 | 3.07771 | 2.38049 | 4.27600 |
| **2** | 4.73160 | 54.12430 | 13.31282 | 8.60650 | 5.80710 | 13.41768 |
| **3** | 2.75465 | 67.22043 | 15.94346 | 8.71049 | 4.40773 | 0.96324 |
| **4** | 2.43090 | 37.06928 | 55.94196 | 1.51929 | 1.51929 | 1.51929 |
| **5** | 2.61471 | 96.84637 | 0.13473 | 0.13473 | 0.13473 | 0.13473 |
| **6** | 0.09339 | 99.53306 | 0.09339 | 0.09339 | 0.09339 | 0.09339 |
| **7** | 0.06989 | 99.65058 | 0.06988 | 0.06988 | 0.06988 | 0.06988 |
| **8** | 0.01879 | 99.90644 | 0.01869 | 0.01869 | 0.01869 | 0.01869 |
| **9** | 0.10197 | 99.51720 | 0.09521 | 0.09521 | 0.09521 | 0.09521 |
| **10** | 0.41072 | 98.84858 | 0.18517 | 0.18517 | 0.18517 | 0.18517 |
| **11** | 1.68349 | 93.51384 | 1.20067 | 1.20067 | 1.20067 | 1.20067 |
| **12** | 2.70069 | 96.70133 | 0.14950 | 0.14950 | 0.14950 | 0.14950 |
| **13** | 49.59636 | 44.56774 | 1.45898 | 1.45898 | 1.45898 | 1.45898 |
| **14** | 99.33753 | 0.13249 | 0.13249 | 0.13249 | 0.13249 | 0.13249 |
| **15** | 3.93150 | 93.59238 | 0.61903 | 0.61903 | 0.61903 | 0.61903 |
| **16** | 33.93239 | 60.39669 | 1.41773 | 1.41773 | 1.41773 | 1.41773 |
| **17** | 0.02666 | 99.86670 | 0.02666 | 0.02666 | 0.02666 | 0.02666 |
| **18** | 0.00036 | 99.99825 | 0.00035 | 0.00035 | 0.00035 | 0.00035 |
| **19** | 0.01995 | 99.90026 | 0.01995 | 0.01995 | 0.01995 | 0.01995 |
| **20** | 0.02314 | 99.88669 | 0.02254 | 0.02254 | 0.02254 | 0.02254 |
| **21** | 0.03592 | 99.90613 | 0.01449 | 0.01449 | 0.01449 | 0.01449 |
| **22** | 0.13499 | 99.32504 | 0.13499 | 0.13499 | 0.13499 | 0.13499 |
| **23** | 0.75202 | 98.93517 | 0.07820 | 0.07820 | 0.07820 | 0.07820 |
| **24** | 15.48279 | 81.64149 | 0.71893 | 0.71893 | 0.71893 | 0.71893 |
| **25** | 0.00236 | 99.98822 | 0.00236 | 0.00236 | 0.00236 | 0.00236 |
| **26** | 0.00000 | 100.00000 | 0.00000 | 0.00000 | 0.00000 | 0.00000 |
| **27** | 0.03032 | 99.84841 | 0.03032 | 0.03032 | 0.03032 | 0.03032 |
| **28** | 0.00000 | 100.00000 | 0.00000 | 0.00000 | 0.00000 | 0.00000 |
| **29** | 0.00327 | 99.98367 | 0.00327 | 0.00327 | 0.00327 | 0.00327 |
| **30** | 0.08158 | 99.59211 | 0.08158 | 0.08158 | 0.08158 | 0.08158 |
| **31** | 0.09025 | 99.54875 | 0.09025 | 0.09025 | 0.09025 | 0.09025 |
| **32** | 49.10495 | 45.36182 | 1.38331 | 1.38331 | 1.38331 | 1.38331 |
| **33** | 98.23178 | 0.35364 | 0.35364 | 0.35364 | 0.35364 | 0.35364 |
| **34** | 0.21781 | 0.21781 | 98.91097 | 0.21781 | 0.21781 | 0.21781 |
| **35** | 1.18453 | 1.18453 | 94.07734 | 1.18453 | 1.18453 | 1.18453 |
| **36** | 0.02060 | 0.02060 | 99.89701 | 0.02060 | 0.02060 | 0.02060 |
| **37** | 8.74315 | 26.73696 | 3.01433 | 39.33044 | 19.16079 | 3.01433 |
| **38** | 0.94837 | 57.06267 | 0.94837 | 0.94837 | 39.14385 | 0.94837 |
| **39** | 0.34851 | 98.25744 | 0.34851 | 0.34851 | 0.34851 | 0.34851 |
| **40** | 14.12021 | 2.71176 | 2.71147 | 75.03362 | 2.71147 | 2.71147 |
| **41** | 0.59931 | 0.59931 | 0.59931 | 97.00347 | 0.59931 | 0.59931 |
| **42** | 99.99688 | 0.00269 | 0.00011 | 0.00011 | 0.00011 | 0.00011 |
| **43** | 0.96260 | 6.11003 | 0.96385 | 0.96260 | 0.96260 | 90.03833 |
| **44** | 0.00354 | 0.00354 | 0.01589 | 0.00354 | 0.00354 | 99.96996 |
| **45** | 0.13825 | 0.13825 | 3.67758 | 0.13825 | 0.13825 | 95.76940 |
| **46** | 1.00809 | 1.00809 | 38.10212 | 1.00809 | 1.00809 | 57.86553 |
| **47** | 0.16141 | 0.16141 | 0.16141 | 0.16141 | 0.16141 | 99.19296 |
| **48** | 0.00000 | 0.00000 | 0.00000 | 0.00000 | 0.00000 | 100.00000 |
| **49** | 0.00022 | 0.00001 | 0.00019 | 0.00001 | 0.00001 | 99.99957 |
| **50** | 42.17821 | 0.65379 | 36.00700 | 0.64936 | 0.64936 | 19.86228 |
| **51** | 0.49497 | 0.50283 | 63.35818 | 0.49497 | 0.49497 | 34.65409 |
| **52** | 0.10403 | 0.10403 | 99.47985 | 0.10403 | 0.10403 | 0.10403 |
| **53** | 0.00761 | 0.00761 | 99.96193 | 0.00761 | 0.00761 | 0.00761 |
| **54** | 0.00000 | 0.00001 | 99.99999 | 0.00000 | 0.00000 | 0.00000 |
| **55** | 0.59901 | 8.19844 | 89.40554 | 0.59901 | 0.59901 | 0.59901 |
| **56** | 0.01340 | 0.01340 | 99.93299 | 0.01340 | 0.01340 | 0.01340 |

| | | | | | | |
|---|---|---|---|---|---|---|
| **57** | 0.02205 | 0.04502 | 0.11818 | 0.02205 | 0.02205 | 99.77064 |
| **58** | 0.28989 | 0.28989 | 8.10116 | 0.28989 | 0.28989 | 90.73929 |
| **59** | 0.00799 | 0.00799 | 0.00799 | 0.00799 | 0.00799 | 99.96004 |
| **60** | 0.00000 | 0.00000 | 0.00000 | 0.00000 | 0.00000 | 99.99999 |
| **61** | 0.00534 | 0.00534 | 0.00534 | 0.00534 | 0.00534 | 99.97328 |
| **62** | 0.00000 | 0.00000 | 0.00000 | 0.00000 | 0.00000 | 100.00000 |
| **63** | 0.08302 | 1.26181 | 0.08302 | 0.08302 | 0.08302 | 98.40613 |
| **64** | 2.33239 | 51.22574 | 2.33239 | 2.33239 | 2.33239 | 39.44468 |
| **65** | 0.07569 | 0.07569 | 0.07569 | 0.07569 | 0.07569 | 99.62157 |
| **66** | 0.01460 | 0.01460 | 0.01460 | 0.01460 | 0.01460 | 99.92700 |
| **67** | 0.00969 | 0.00969 | 0.00969 | 0.00969 | 0.00969 | 99.95157 |
| **68** | 0.05029 | 0.05029 | 0.05029 | 0.05029 | 0.05029 | 99.74854 |
| **69** | 0.01089 | 0.01089 | 0.01089 | 0.01089 | 0.01089 | 99.94557 |
| **70** | 0.01658 | 0.01658 | 0.01658 | 0.01658 | 0.01658 | 99.91712 |
| **71** | 0.09280 | 0.09280 | 0.09280 | 0.09280 | 0.09280 | 99.53601 |
| **72** | 0.01062 | 0.01062 | 0.01062 | 0.01062 | 0.01062 | 99.94691 |
| **73** | 0.00000 | 0.00000 | 0.00000 | 0.00000 | 0.00000 | 100.00000 |
| **74** | 0.01570 | 0.01570 | 0.01570 | 0.01570 | 0.01570 | 99.92151 |
| **75** | 0.00000 | 0.00000 | 0.00000 | 0.00000 | 0.00000 | 99.99999 |
| **76** | 3.39975 | 37.67494 | 3.39975 | 3.39975 | 3.39975 | 48.72606 |
| **77** | 0.69751 | 0.69751 | 0.69751 | 0.69751 | 0.69751 | 96.51246 |

**Table S4.** Predicted $\lambda_{max}$ based on pure $A_1$ records and nomogram fitting

| Species | Photoreceptor type | | | | | | |
|---|---|---|---|---|---|---|---|
| | RH1 | SWS2 | RH2 | RH2/ LWS2 | LWS2 | LWS2/ LWS1 | LWS1 |
| Curimatidae | | | | | | | |
| *C. magdalenae* | | | | | | | |
| Reference $\lambda_{maxA1}$ | 504 | 442 | 475 | 514 | 530 | 544 | 560 |
| Reference $\lambda_{maxA2}$ | 536 | 455 | 496 | 550 | 573 | 595 | 622 |
| Erythrinidae | | | | | | | |
| *H. microlepis* | | | | | | | |
| Reference $\lambda_{maxA1}$ | 503 | — | 478 | 514 | 530 | 544 | 560 |
| Reference $\lambda_{maxA2}$ | 534 | — | 500 | 550 | 574 | 596 | 622 |
| Bryconidae | | | | | | | |
| *B. chagrensis* | | | | | | | |
| Reference $\lambda_{maxA1}$ | 504 | 446 | 472 | 514 | 530 | — | 560 |
| Reference $\lambda_{maxA2}$ | 536 | 460 | 492 | 550 | 574 | — | 622 |
| Characidae | | | | | | | |
| *G.atracaudatus* | | | | | | | |
| Reference $\lambda_{maxA1}$ | 504 | 440 | 480 | 514 | 530 | — | — |
| Reference $\lambda_{maxA2}$ | 536 | 453 | 503 | 550 | 574 | — | — |
| *B. gonzalezi* | | | | | | | |
| Reference $\lambda_{maxA1}$ | 504 | 449 | 479 | 514 | 530 | 544 | 560 |
| Reference $\lambda_{maxA2}$ | 536 | 464 | 511 | 550 | 574 | 596 | 622 |
| *R. guatemalensis* | | | | | | | |
| Reference $\lambda_{maxA1}$ | 502 | 447 | 480 | — | 530 | — | — |
| Reference $\lambda_{maxA2}$ | 533 | 462 | 503 | — | 574 | — | — |
| *A. ruberrimus* | | | | | | | |
| Reference $\lambda_{maxA1}$ | 504 | 447 | 472 | 514 | 529 | — | — |
| Reference $\lambda_{maxA2}$ | 536 | 462 | 492 | 550 | 572 | — | — |

**Table S5.** Kruskal Wallis Rank Sum Test of photoreceptors $\lambda_{max}$ across species.

| Photoreceptor type | |
|---|---|
| Rods | chi-squared = 105, df = 6, p-value < 2.2e-16 |
| Long-green cones | chi-squared = 50.428, df = 6, p-value = 3.859e-09 |
| Blue cones | chi-squared = 19.53, df = 5, p-value = 0.001531 |

**Table S6.** Pair Wise Wilcoxon Rank Sum Test

**Rods**

| | *A. ruberrimus* | *B. chagrensis* | *B. gonzalezi* | *C. magdalenae* | *G. atracaudatus* | *H. microlepis* |
|---|---|---|---|---|---|---|
| *B. chagrensis* | **0.00022** | - | - | - | - | - |
| *B. gonzalezi* | 0.86445 | **1.90E-05** | - | - | - | - |
| *C. magdalenae* | **1.10E-08** | **1.10E-05** | **1.10E-11** | - | - | - |
| *G. atracaudatus* | 0.23285 | **0.0104** | 0.34286 | **9.10E-08** | - | - |
| *H. microlepis* | **4.00E-06** | 0.16282 | **1.80E-07** | **0.00044** | **0.00029** | - |
| *R. guatemalensis* | 0.8725 | **2.90E-05** | 0.50535 | **3.10E-10** | 0.16282 | **7.80E-07** |

**Long-green cones**

| | *A. ruberrimus* | *B. chagrensis* | *B. gonzalezi* | *C. magdalenae* | *G. atracaudatus* | *H. microlepis* |
|---|---|---|---|---|---|---|
| *B. chagrensis* | **0.03097** | - | - | - | - | - |
| *B. gonzalezi* | 0.79622 | **0.0032** | - | - | - | - |
| *C. magdalenae* | 0.1089 | 0.1906 | **0.03097** | - | - | - |
| *G. atracaudatus* | 0.33539 | **0.00083** | 0.79622 | **0.08762** | - | - |
| *H. microlepis* | **0.03097** | 0.79622 | **0.00355** | 0.06886 | **0.00083** | - |
| *R. guatemalensis* | 0.79622 | **3.40E-05** | 0.25445 | **0.02016** | 0.25445 | **7.30E-05** |

**Blue cones**

| | *A. ruberrimus* | *B. chagrensis* | *B. gonzalezi* | *C. magdalenae* | *G. atracaudatus* |
|---|---|---|---|---|---|
| *B. chagrensis* | 0.5948 | - | - | - | - |
| *B. gonzalezi* | 0.5221 | 0.3559 | - | - | - |
| *C. magdalenae* | 0.3559 | 0.1896 | 0.6065 | - | - |
| *G. atracaudatus* | **0.0082** | **0.0082** | **0.0497** | 0.3559 | - |
| *R. guatemalensis* | 0.6065 | 0.7561 | 0.5221 | 0.1896 | **0.023** |

*Shaded boxes denote species collected in murky environments. Blue cones were not found in *H. microlepis*.

**Table S7.** Kruskal Wallis Rank Sum Test of rods $\lambda_{max}$ among individuals

| Species | |
|---|---|
| *A. ruberrimus* | chi-squared = 8.0748, df = 10, p-value = 0.6215 |
| *B. chagrensis* | chi-squared = 27.408, df = 18, p-value = 0.07166 |
| *B. gonzalezi* | chi-squared = 7.5498, df = 11, p-value = 0.753 |
| *C. magdalenae* | chi-squared = 6.4917, df = 10, p-value = 0.7724 |
| *G. atracaudatus* | chi-squared = 13.09, df = 10, p-value = 0.2187 |
| *H. microlepis* | chi-squared = 14.399, df = 12, p-value = 0.2759 |
| *R. guatemalensis* | chi-squared = 9.6662, df = 13, p-value = 0.721 |

**Table S8.** Sampling

| Species | Family | Locality | Province | Coordinates (Lat/Lon) |
|---|---|---|---|---|
| *Bryconamericus gonzalezi* | Characidae | Quebrada Gavilán (Clear-water) | Bocas del Toro | 9.272083, 82.509444 |
| *Roeboides guatemalensis* | Characidae | Rio Juan Grande (Clear-water) | Colon | 9.136361, -79.723576 |
| *Piabucina panamensis* | Lebiasinidae | Rio Juan Grande (Clear-water) | Colon | 9.136361, -79.723580 |
| *Gephyrocarax atracaudatus* | Characidae | Rio Juan Grande (Clear-water) | Colon | 9.136361, -79.723581 |
| *Hyphessobrycon panamensis* | Characidae | Rio Juan Grande (Clear-water) | Colon | 9.136361, -79.723587 |
| *Bryconamericus emperador* | Characidae | Rio Juan Grande (Clear-water) | Colon | 9.136361, -79.723594 |
| *Astyanax ruberrimus* | Characidae | Rio Juan Grande (Clear-water) | Colon | 9.136361, -79.723598 |
| *Hoplias microlepis* | Erythrinidae | Gamboa Dock (Murky-water) | Colon | 9.113547, -79.691149 |
| *Brycon chagrensis* | Bryconidae | Isla de los Monos (Murky-water) | Colon | 9.119149, -79.779212 |
| *Cyphocharax magdalenae* | Curimatidae | Los Canelos (Murky-water) | Veraguas | 8.09508, 80.64141 |
| *Carnegiella strigata** | Gasteropelecidae | Coropina creek, Berlijn (Black-water) | Para | 5.397194, 55.184306 |
| *Serrasalmus rhombeus** | Serrasalmidae | Van-Blommenstein Lake (Clear-water) | Brokopondo | 4.960222, 54.982444 |
| *Crenuchus spilurus** | Crenuchidae | Coropina creek, Berlijn (Black-water) | Para | 5.397194 55.184306 |

*Denotes samples collected in Suriname where only two transcriptomes per species were performed

**Table S9.** Evolutionary models

| Opsin | Model | Data |
|---|---|---|
| SWS | LG+I+G+F | AA |
| RH2-RH1 | LG+G+F | AA |
| LWS | LG+I+G | AA |
| Multilocus phylogeny* | GTR+I+G | DNA |

* GTR+I+G was used for each partition for every codon position.

**Table S10.** Combinations of the occurrence of tuning sites

| | Tuning sites | | |
|---|---|---|---|
| **Combination** | **A164S** | **F261Y** | **A269T** |
| 1 | A | F | A |
| 2 | S | Y | T |
| 3 | A | Y | A |
| 4 | A | Y | T |
| 5 | S | F | A |
| 6 | S | Y | A |
| 7 | A | F | T |

### Supplementary Figures

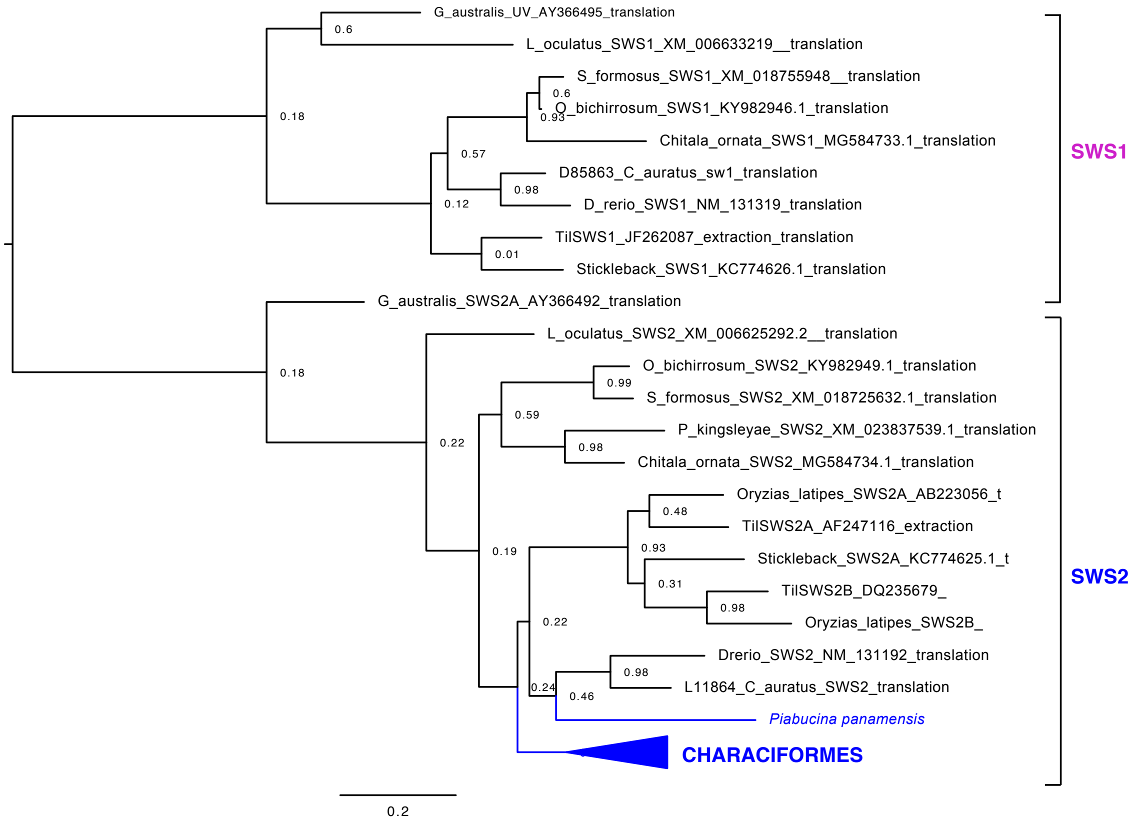


**Figure S1. SWS opsin tree of Characiformes.** SWS opsin maximum-likelihood phylogenetic tree based on amino-acid sequences of Characiformes, Osteoglossiformes, *Geotria australis* (lamprey), *Lepisosteus oculatus* (Spotted gar), *Oryzias latipes* (medaka), *Gasterosteus aculeatus* (stickleback), *Carassius auratus* (goldfish), and *Danio rerio* (zebrafish). Bootstrap support over 75% is shown. This tree confirms that SWS1 seems to be lost in Characiformes. Characiform species are represented as the compressed blue-colored clade although *P. panamensis* clusters outside the rest of Characiformes.

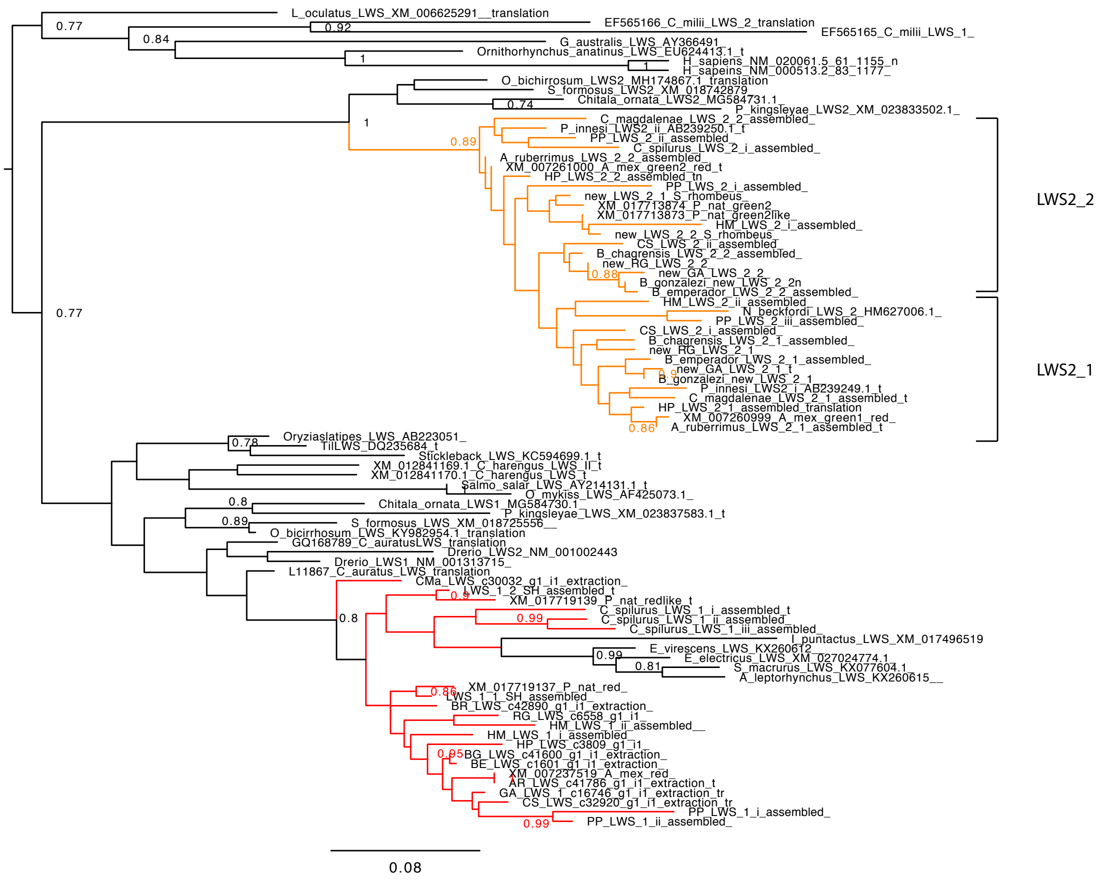

**Figure S2. LWS opsin tree of Characiformes.** LWS opsin maximum-likelihood phylogenetic tree based on LWS-amino-acid sequences of Characiformes, Osteoglossiformes, Siluriformes, Gymnotiformes, *Geotria australis* (lamprey), *Ornithorhynchus anatinus* (platypus), *Homo sapiens* (humans), *Callorhinchus milii* (Elephant shark), *Lepisosteus oculatus* (Spotted gar), *Oryzias latipes* (medaka), *Gasterosteus aculeatus* (stickleback), *Clupea harengus* (herring), *Salmo salar* (salmon), *Onchrynchus mykiss* (trout), *Carassius auratus* (goldfish), and *Danio rerio* (zebrafish). Bootstrap support over 75% is shown. This tree confirms that LWS1 and LWS2 arose after the divergence of the spotted gar, probably as a product of TGD. Notice the clustering of characiform LWS2 opsins with the osteoglossimorph LWS2 opsins, and the clustering of Siluriformes and Gymnotiformes within the characiform LWS1 opsin clade. Characiform species are color coded for LWS2 in orange and LWS1in red.

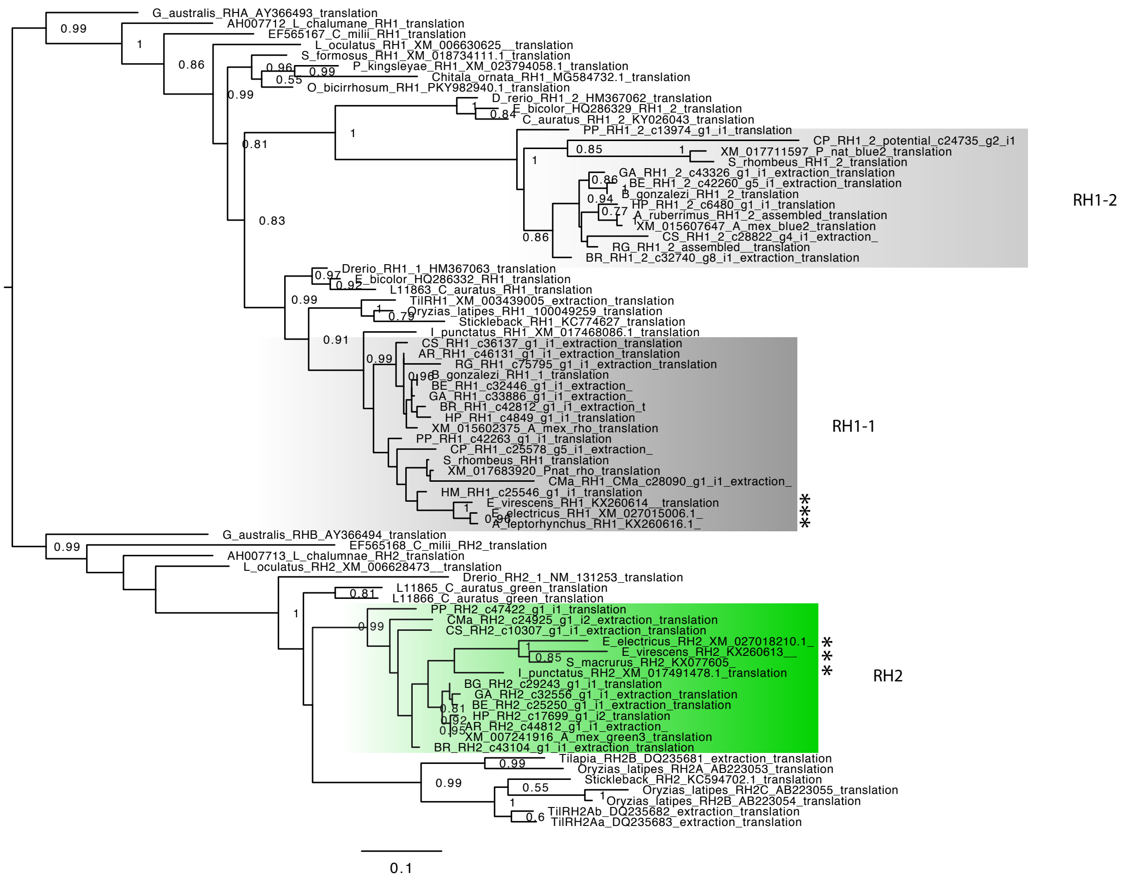


**Figure S3. RH1-RH2 opsin tree of Characiformes.** RH1-RH2 opsin maximum-likelihood phylogenetic tree based on RH1-RH2-amino-acid sequences of Characiformes, Osteoglossiformes, Siluriformes, Gymnotiformes, Cypriniformes, *Geotria australis* (lamprey), *Latimeria calumnae* (coelacant), *Callorhinchus milii* (Elephant shark), *Lepisosteus oculatus* (Spotted gar), *Oryzias latipes* (medaka), *Gasterosteus aculeatus* (stickleback). Bootstrap support over 75% is shown. This tree confirms that RH1-2 arose after the divergence of the spotted gar, probably as a product of TGD. Notice the clustering of characiform RH1-2 opsins with the cyprinimorphs surviving RH1-2 opsins. Characiform species clades are shown in different colors (*RH1-2 in gray, RH1-1 in black, and RH2 in green*). * denotes siluriform species nested within characiform clades.

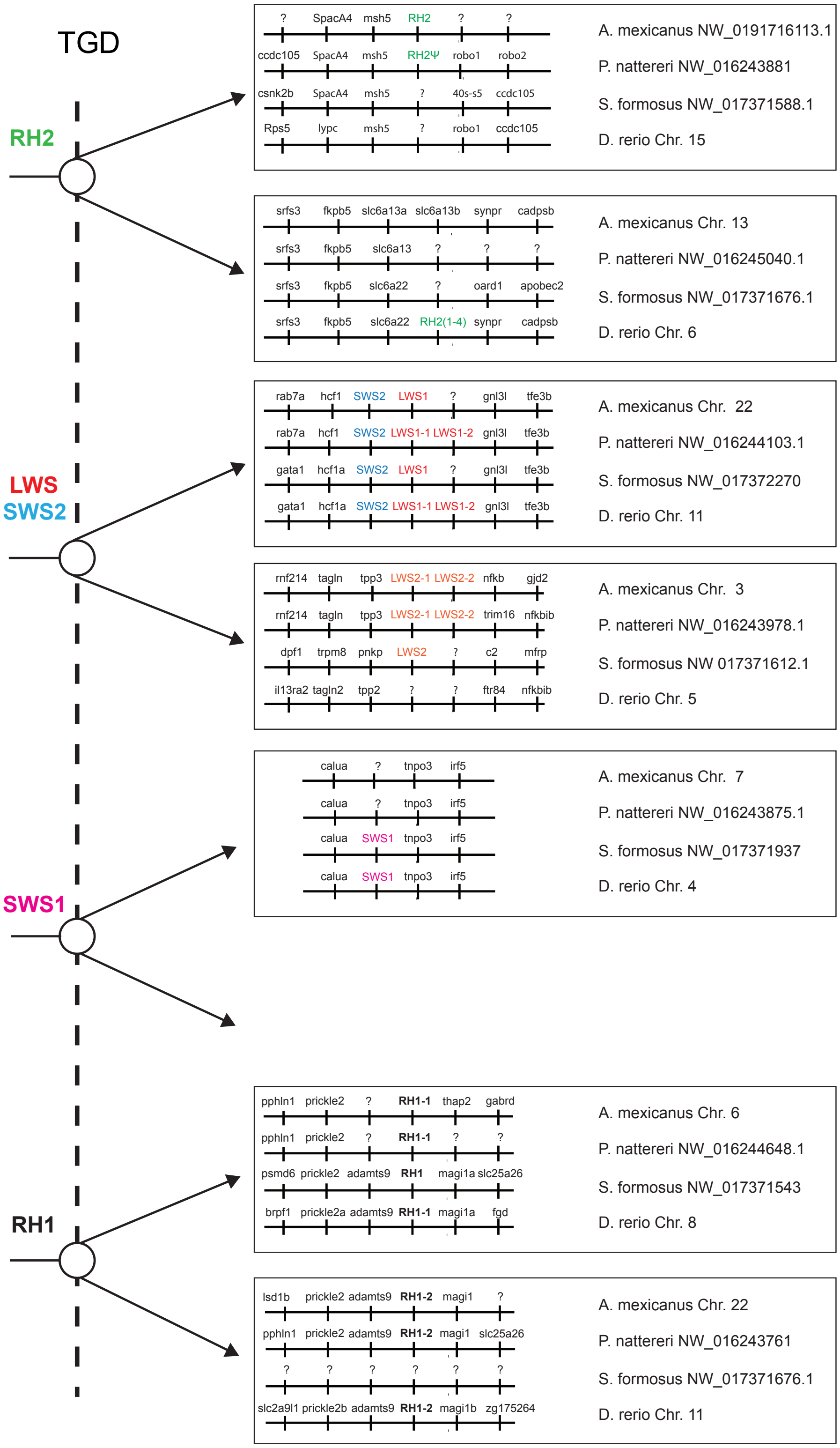


**Figure S4.** Scenario for opsins duplications produced by TGD and based on synteny analysis of putative chromosomal regions surrounding different opsins in *Danio rerio, Scleropages formosus, Astyanax mexicanus* and *Pygocentrus nattereri.*

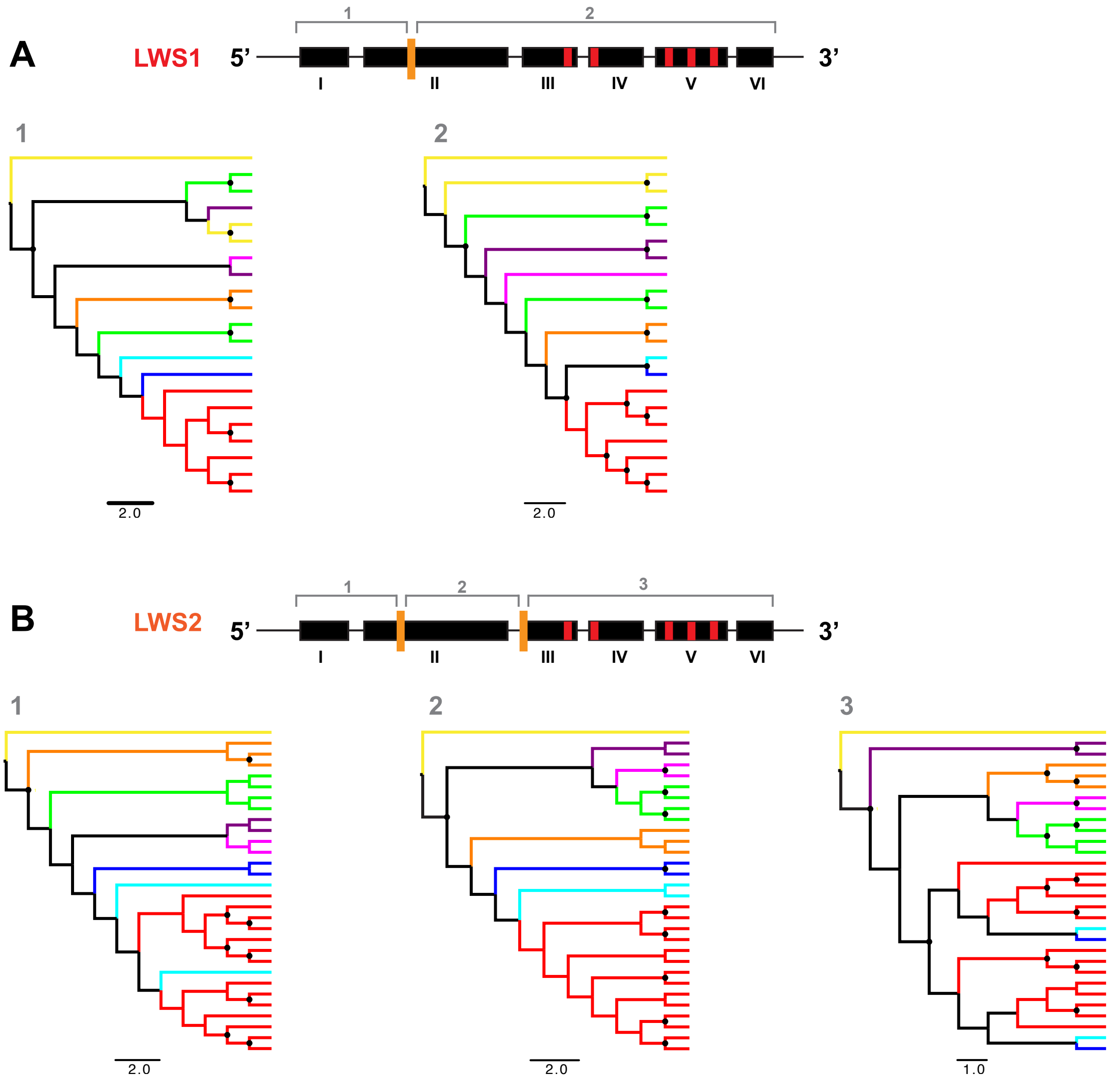


**Figure S5.** Overview of LWS gene conversion analysis. Schematic representation of exon structure of LWS1 and LWS2 is shown, where roman numbers indicate the exon number. Orange vertical bars denote the breakpoints in each LWS opsin and red bars denote the five "key sites" of Yokoyama and Radlwimmer (2001) ). Nucleotide trees based on 15 characiform LWS opsins are shown where each family is color coded *(Crenuchidae in yellow, Lebiasinidae in orange, Serrasalmidae in green, Erythrinidae in purple, Curimatidae in violet, Gasteropelecidae in blue, Bryconidae in light-blue, and Characidae in red)*. In tree #3 for the LWS2, it is suggested that Bryconidae, Gasteropelecidae and Characidae share a LWS2 duplication. For each fragment we used RAXML to build maximum-likelihood trees. We ran 10 searches for the best tree and 1000 bootstrap replicates, performed in RAXML 8.0 on CIPRES. A black circle denotes bootstrap support over 75%.

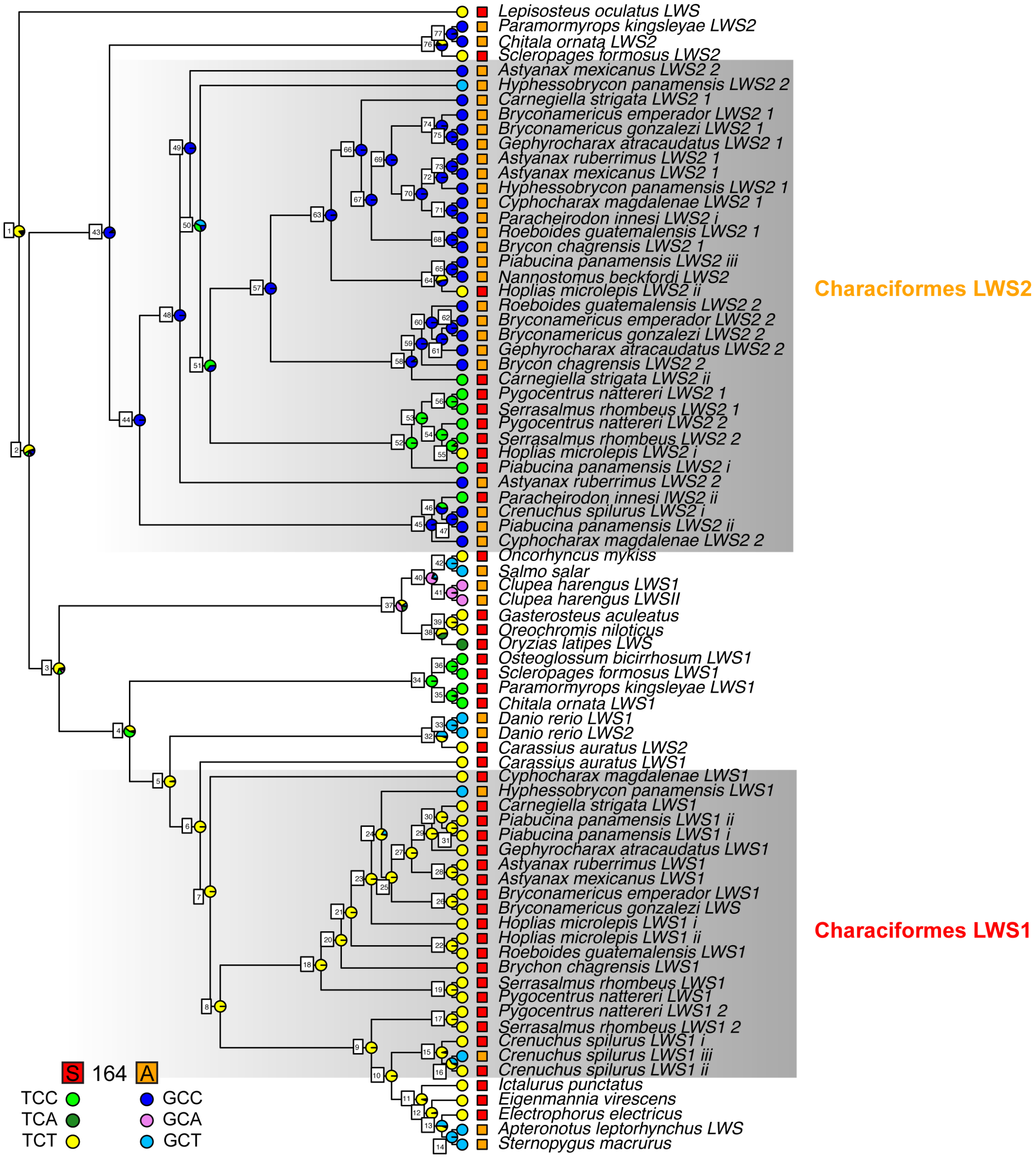


**Figure S6.** Ancestral state reconstruction results for the spectral tuning site 164. Squares indicate whether the opsin gene sequence has alanine or serine. Codons at site 164 are color coded in colored filled circles. Pie charts on the nodes indicate the scaled likelihoods (calculated using the ace function in APE) of each specific combination. Nodes are also labeled as in Table S4.

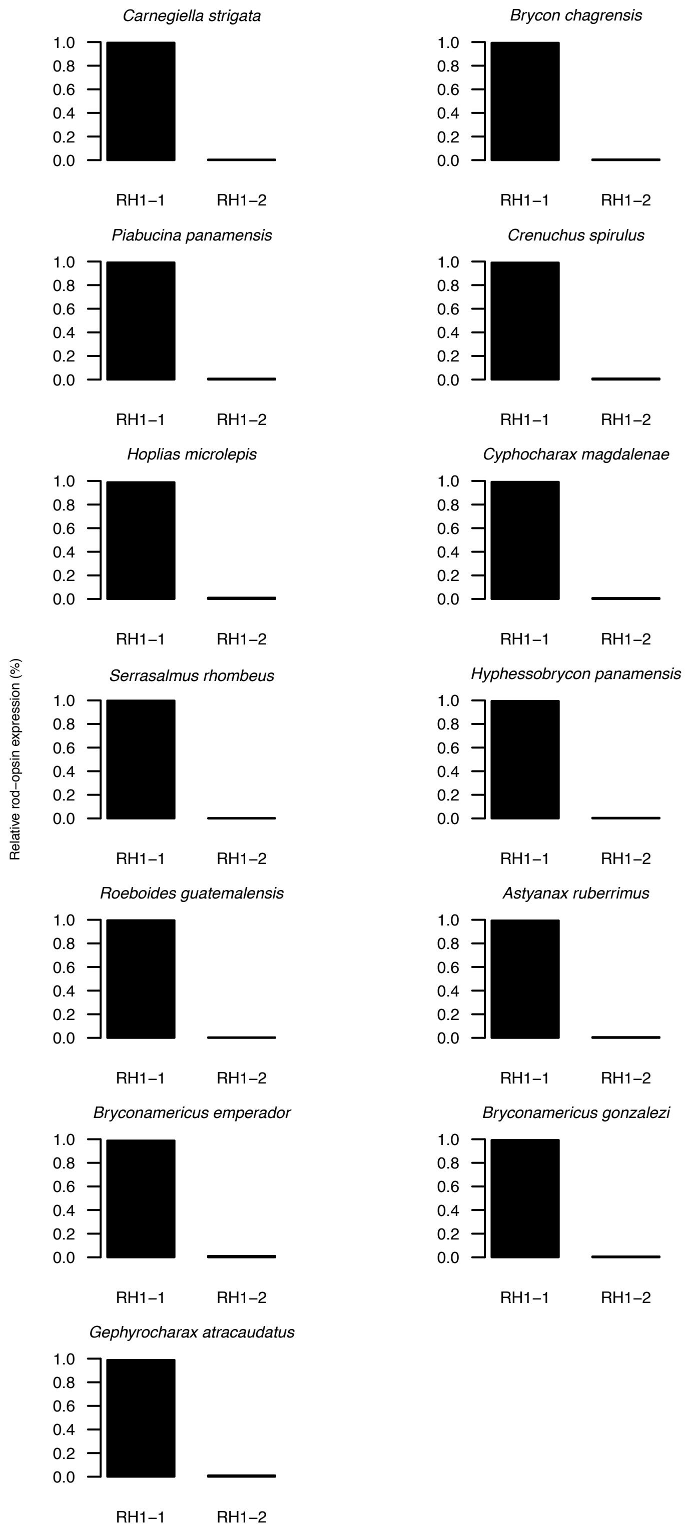


**Figure S7.** Rodopsin expression in the characiform species examined.

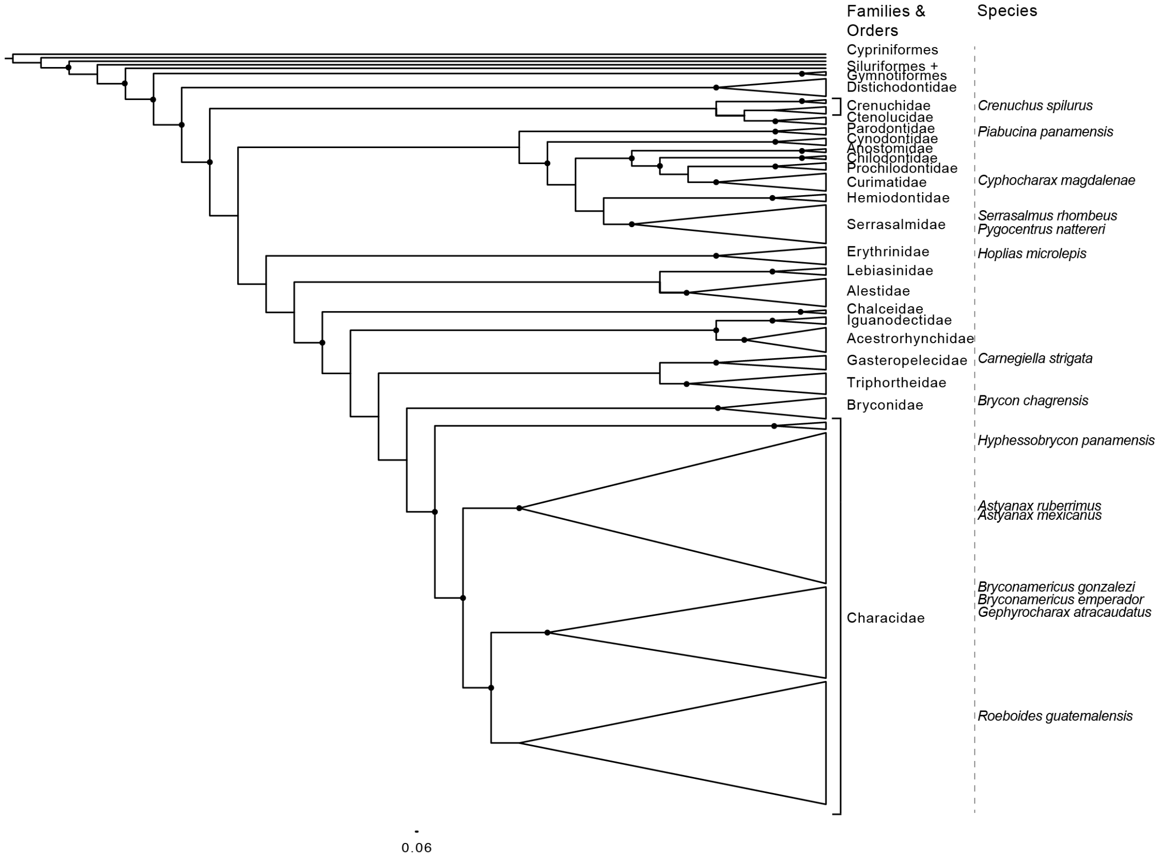


**Figure S8.** Phylogeny of 228 Characiformes based on genomic sequences. Species sampled in Panama and Suriname are included. Filled circles denote bootstrap support over 75%.
